# An agent-based model suggests how senescent cell behavior and matrix mechanics drive pulmonary fibrosis in aged mice

**DOI:** 10.64898/2026.05.01.722023

**Authors:** Mackenzie L. Skelton, Julie Leonard-Duke, Leilani R. Astrab, Joshua A. Goedert, Riley T. Hannan, Shayn M. Peirce, Jeffrey M. Sturek, Steven R. Caliari

## Abstract

Idiopathic pulmonary fibrosis (IPF) is a progressive and ultimately fatal disease of aging, driven by dysregulated fibroblast activation and accompanied by collagen accumulation in the lung interstitium, resulting in tissue stiffening. While the accumulation of senescent cells has been increasingly implicated in IPF pathogenesis, understanding the reciprocal dynamics of senescent fibroblast levels and evolving tissue mechanics is difficult to achieve with experimental approaches alone. To address this limitation, we developed an agent-based model (ABM) of fibroblast activation in the lung that couples cell behavior to the dynamic mechanical changes accompanying fibrosis. This model was parameterized entirely from experimental data in young mice to enable robust validation and then adapted to fit aged mouse biology for additional validation. Both young and aged models accurately reflected changes in collagen accumulation and stiffness burden of experimental systems. We then incorporated senescent cell behavior into the aged model to investigate how senescent cell burden influences fibrosis progression and how cell-cell interactions drive senescent cell accumulation. These simulations identified a unique role for juxtacrine-mediated contact between non-senescent and senescent fibroblasts in expanding the total senescent cell burden. Our ABM also revealed that the timing of immune-mediated senescent cell clearance critically regulates fibrotic outcomes. Together, this ABM provides useful insights into how the interrelated dynamics of tissue mechanics and senescent fibroblasts drive fibrosis progression.

## Introduction

Idiopathic lung fibrosis (IPF) is a devastating condition of pathological scarring in the lung interstitium that hinders breathing. The median survival time for IPF patients from diagnosis is 2-5 years [1,2], and the currently approved therapeutics are not able to stop the progression of the disease. This lack of effective therapeutic options is due in part to IPF’s unknown etiology, highlighting a need to better understand the mechanisms initiating and driving this disease. The drastic increase in IPF prevalence in people aged 65 and above points to a potential link between aging and IPF onset [3–5]. Various aging mechanisms have been implicated in IPF progression such as changes in the aged lung composition [6,7], chronic low-grade inflammation [8], and the accumulation of senescent cells [9–11]. Yet, how these characteristics work in tandem toward the progression of fibrosis remains unclear. Understanding the connection between aging and IPF is crucial for advancing treatment options.

One hallmark of aging is the accumulation of senescent cells, observed across many tissue types [12–14]. Senescence is a state of stable cell cycle arrest where cells exhibit a robust secretory profile, known as the senescence-associated secretory phenotype (SASP) [15,16]. Cellular senescence represents a nexus of pro-inflammatory and pro-fibrotic signaling, through the recruitment of immune cells, the activation of pro-fibrotic myofibroblasts, and the propagation of this damage-induced phenotype [10,11]. Work implicating senescent cells’ influence on fibrosis progression has led to multiple ongoing clinical trials targeting senescent cell clearance [17,18]. Senescent fibroblasts in particular have been shown to exhibit a highly pro-fibrotic SASP [9], and the selective elimination of senescent fibroblasts alone is able to reverse persistent fibrosis in mice [19]. However, senescent cells can also play a beneficial role in tissue repair, and eliminating them completely could lead to adverse effects [20]. Thus, there is a great need to understand what aspects of senescent fibroblast behavior contribute most to fibrosis progression.

Fibroblasts are the primary effector cell of scar tissue formation through the deposition and contraction of extracellular matrix (ECM) [21,22]. In the presence of mechanical and biochemical pro-fibrotic cues, quiescent fibroblasts activate into myofibroblasts of distinct subtypes that accomplish these functions, discussed here as proto-myofibroblasts and contractile myofibroblasts [23,24]. While senescent fibroblasts have been identified in fibrotic tissue, their origin and dynamics are largely unknown. Some have shown that fully activated, contractile myofibroblasts, may trigger senescence to avert apoptosis [25], but the relative influence of this pathway on fibrosis progression has not been studied. Additionally, the contribution of senescent fibroblasts to fibrosis progression relative to other myofibroblasts is not known. These questions warrant a systematic study that investigates the dynamics of these key fibroblast subtypes in fibrosis progression.

Extracellular matrix (ECM) cues have been long recognized for their contribution to fibrosis [26–28]. Specifically, the increase in matrix stiffness caused by activated fibroblasts depositing excess ECM drives a potent positive feedback loop promoting fibrosis progression [24,29,30]. More recent studies have also highlighted the importance of viscoelasticity, or the ability of a tissue to relax stress, in preventing fibroblast activation [31–34]. These complex changes in ECM mechanics occur not only with fibrosis progression, but also with age in the absence of fibrosis [35–37]. The ECM of many tissues becomes stiffer with age [38–40], but studies have shown conflicting and tissue-specific information regarding the changes in viscoelastic character with age [41–46]. The pivotal role of viscoelastic mechanics in fibrosis development necessitates their integration in a unified model.

With the dynamic mechanical and cellular complexity comprising the IPF landscape, a computational approach is required for understanding the role of each of these factors in the disease. Agent-based models (ABMs) have served as a powerful tool for understanding cellular interactions driving fibrosis across various organs [47–51]. For IPF specifically, previous models have generated novel insights into the role of intercellular interactions [52,53], the source of pro-fibrotic cytokines [54], tissue mechanics [55], and organization [56] on the progression of the disease. Additionally, a recent model investigating senescent cell behavior in wound healing has provided valuable insights into the temporal influence of senescent cell clearance [57]. However, there is a need to investigate the role of senescent cell dynamics in a tissue-specific model of IPF.

This paper presents a novel ABM of pulmonary fibrosis incorporating the dynamic mechanical changes present in IPF and aging to investigate the role of senescent fibroblast behavior on fibrosis progression. We utilized the bleomycin model of lung fibrosis in young and aged mice, as this is the standard animal model for studying IPF. From these mouse studies we generated mouse lung histology images to specify 2D lung microarchitectures of both young and aged mice in the ABM, and we validated model outputs by comparing predictions to mechanical and biochemical measurements of young and aged mice. Senescent fibroblast behavior was then incorporated into the aged mouse ABM, which subsequently improved validation metrics. Finally, senescent fibroblast behaviors and dynamics were investigated for their influence on fibrosis progression.

## Methods

### Bleomycin mouse model

#### Young and aged bleomycin mouse model

All animal studies were approved by the University of Virginia’s Institutional Animal Care and Use Committee (IACUC). Male C57BL/6J mice were purchased from Jackson Laboratory at 10 weeks old and kept in the vivarium for two weeks before beginning the study. For bleomycin installation of young mice, animals were sedated, and 1 U/kg bleomycin was delivered trans-orally. Mice were monitored and scored daily following bleomycin installation for changes in weight, appearance, activity, posture, and breathing. Animals with severe changes in any of these categories or with an overall health burden that exceeded the levels specified in our IACUC approved protocol were humanely euthanized via ketamine and opening of the thoracic cavity. Mice that maintained appropriate health were given another two rounds of bleomycin, each two weeks apart. Two weeks following the final administration of bleomycin, lungs were harvested for mechanical testing and histological evaluation.

Similarly, aged mouse experiments were performed on mice that were obtained at 8-10 weeks old and aged to 15 months (67 weeks) before bleomycin instillation. For the multi-dose model, animals were sedated, 0.3 U/kg bleomycin was delivered trans-orally, then mice were monitored daily. Aged mice were given a lower dose of bleomycin to prevent high levels of mortality that have been shown for aged mice [58]. Mice that maintained sufficient health metrics were given another two rounds of bleomycin, each two weeks apart. Then, two weeks following the final administration of bleomycin, mice were sacrificed, and lungs were harvested for mechanical testing and histological analysis. For the single dose group, mice were given 1 U/kg bleomycin and lungs were harvested two weeks later.

#### Mechanical characterization of mouse lungs

Mouse lungs were maintained on ice after harvesting and mechanics were tested within 48 hours of harvest. To investigate the mechanics of the interior of the lung lobe, left lungs were dissociated from the remaining lobes, aligned in a rat heart slicer (Zivic Instruments, 0.5 mm spacing) and cut in half along the coronal plane using a razor blade. The lungs were then adhered to a 15 mm petri dish using a silicone-based adhesive with the interior facing upward to allow submersion in PBS. Indentation tests were performed on a Piuma nanoindenter (Optics 11 Life, Netherlands) using a 50 µm glass probe attached to a cantilever with a spring constant of 0.47 N/m. Indentations were done to a depth of 4 µm at a constant time ramp over 2 seconds and repeated across the ranges indicated.

#### Histological characterization of mouse lungs

Following mechanical characterization, mouse lungs were fixed in 4% paraformaldehyde for 20 minutes and then kept in 30% sucrose for a minimum of 24 hours, or until lungs sunk, before cyrosectioning. 5 µm sections were stained with hematoxylin and eosin (H&E) and Masson’s trichome. Young control mouse histology images used for model initialization can be found in **Fig. S1**, and aged control mouse histology images used for aged model initialization can be found in **Fig. S2**.

### Young Mouse Fibrosis ABM Development

#### Model overview

We utilized NetLogo (version 6.2.2) to develop a 2-dimensional ABM representing a 610 µm x 450 µm area of a lung slice. The ABM’s code is available for download at the Caliari Lab GitHub page (https://github.com/Caliari-Lab/2025_Skelton_SenescenceABM.git). To build this ABM, we leveraged a data-centric approach in which model rules were designed to suit available data. Model behavior focused primarily on behavior of fibroblasts in response to mechanical cues and pro-fibrotic TGFβ1 in the lung from decades of research. Parameters were gathered from both previous ABMs of fibroblast behavior in lung fibrosis [47,54], as well as over 50 published studies. All rules and parameters were based on either direct experimental evidence or extrapolated from experimental data. Details on the origin of all rules and parameters can be found in **Tables 1-9**. A schematic overview of the model workflow and differentiation is shown in **Fig. 1**.

**Figure 1:**
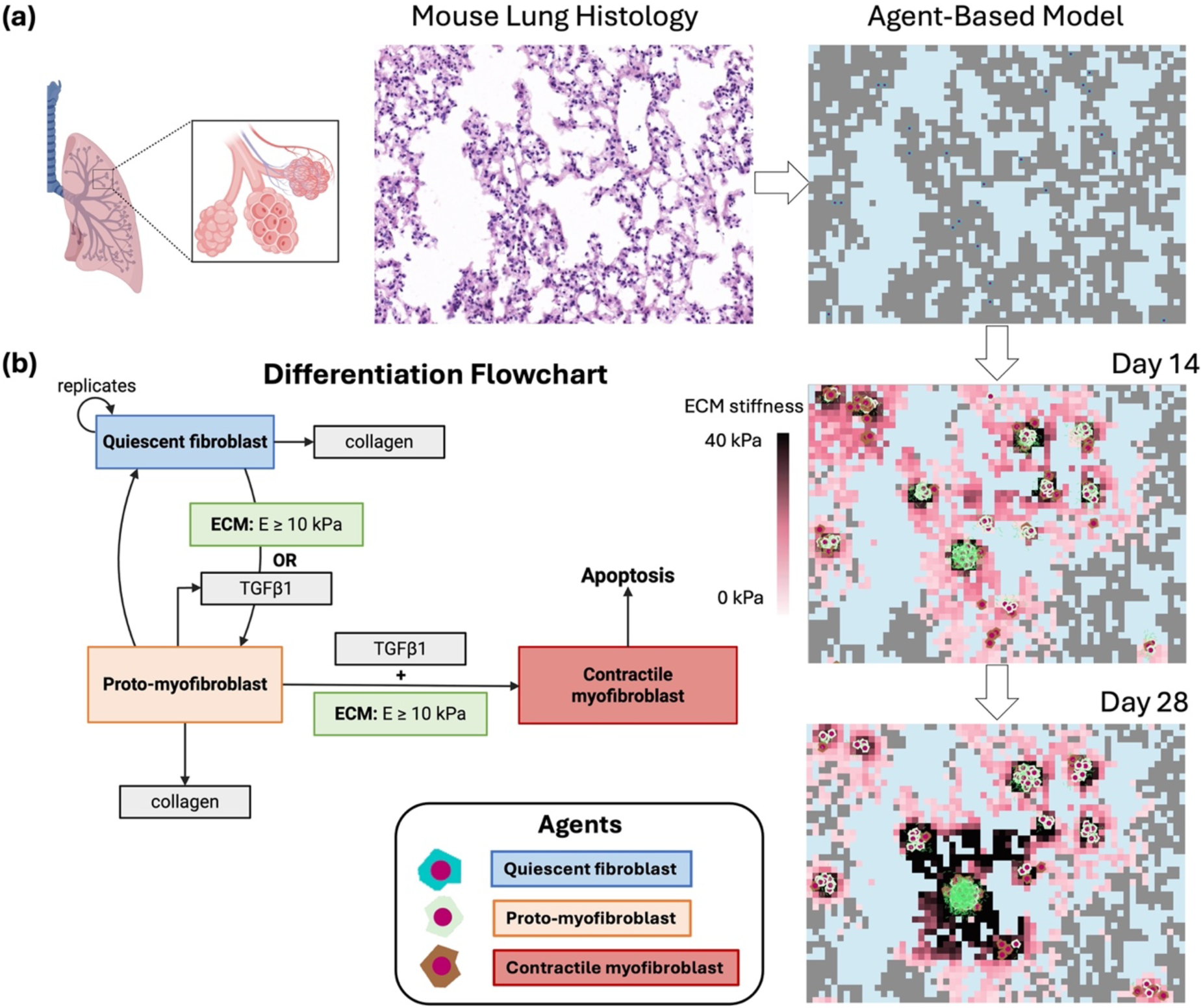
Overview of young mouse ABM design. **(a)** The model depicts alveolar architecture where geometry is defined via imported mouse lung histology images. Agents are initialized in the interstitial space between alveoli and allowed to progress according to the differentiation flowchart depicted in **(b)**. Model initialization begins with fibroblasts and TGFβ1 present in the interstitial space, and the fibroblasts then replicate and differentiate into proto-myofibroblasts. If pro-fibrotic cues persist, proto-myofibroblasts differentiate to contractile myofibroblasts that undergo apoptosis at the end of their lifespan. The model depicts the emergent behavior of fibroblast-rich areas with high stiffness (represented by dark patch colors) and high levels of collagen deposition (represented in green) evident at 28 days.

**Table 1:**
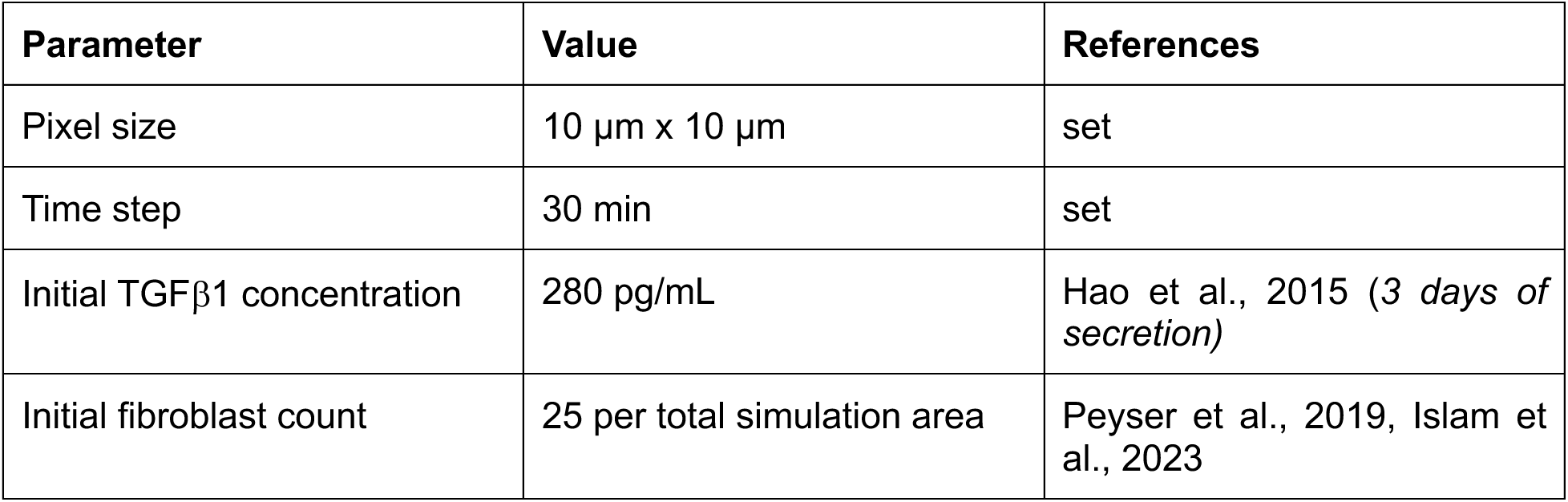
Simulation initialization parameters.

#### Young mouse model initialization and agent definition overview

To initialize the model, we utilized H&E-stained histology images obtained from our *in vivo* mouse studies, described above, to define the architecture of the interstitial space. Full histology images were cropped to the model dimensions (610 x 450 µm), and two crops per histology image were used (**Fig. S1**). Each pixel in the model represented 10 µm, leading to a model space that was 61 x 45 pixels. We assume the fibrotic cascade begins with a damage-causing event that triggers M2-like macrophages to release TGFβ1 [59–63]. At time zero, quiescent fibroblasts and TGFβ1 agents are randomly distributed within the interstitial spaces of the simulation area. Quiescent fibroblasts transition into proto-myofibroblasts, which can either revert to quiescent fibroblasts or transition further into contractile myofibroblasts [64,65]. ECM stiffness and consumption of a threshold of TGFβ1 are required to activate fibroblasts along this lineage, either in an “and” or “or” manner. Quiescent and proto-myofibroblasts deposit collagen into the interstitial space which increases ECM stiffness [66]. Contractile myofibroblasts exert tension on the ECM, which increases the stiffness through strain stiffening behavior [67,68]. A random selection of collagen is degraded each day to simulate the effects of matrix metalloproteinases (MMPs), although the enzymes are not explicitly represented in the model [69]. Proto-myofibroblasts that don’t revert to quiescence or further differentiate and contractile myofibroblasts both die by apoptosis after their designated lifespan. Each time step in the model represents 30 minutes, and the model was run for a range of time representing 7 days to 28 days (336 to 1344 time steps).

#### ECM mechanical behavior

ECM stiffness serves as a primary cue to activate fibroblasts along the myofibroblast lineage [70]. For the initialization of the model, all areas of interstitial space were given a random stiffness value between 1 and 3 kPa, as this is what we and others have measured as the range of stiffness for healthy lung tissue for young mice [46,71]. As fibroblasts deposit collagen, the collagen stiffens the matrix in a manner that is linearly proportional to collagen concentration. This was determined by plotting two relevant datasets [72,73], and the linear model fit was used to determine the extent to which the deposition of collagen by fibroblasts in the model stiffens the matrix. The equation for the line of best fit was then simplified for implementation in the model, and both model fits are shown in **Fig. S3**.

Additionally, the mechanics of the ECM are mediated by strain stiffening and stress relaxation behavior. Contractile myofibroblasts exert high levels of contractile forces on the ECM within a radius of 100 µm, resulting in strain stiffening due to the non-linear elasticity of the fibrous ECM [67,74,75]. The extent of strain stiffening depends on the mechanics of the underlying matrix [76]. Thus, we fit a collection of traction force microscopy data collected from fibroblasts on substrates of various stiffnesses to a logarithmic model, in which there is an inherent capacity for contractile stiffening [77,78]. The best fit model was again simplified for implementation in the model, and both model fits are shown in **Fig. S4**. Strain-induced matrix stiffening was calculated per cell and summed over the total number of contractile cells in each pixel. This strain-induced matrix stiffening was eliminated when contractile myofibroblasts died.

Stress relaxation behavior is another key component of biological tissues [79,80]. Under constant strains, the stress increase in tissues is quickly relaxed by the rearranging of fibers in tissues [81,82]. However, in tissues cells exhibit dynamic strains in response to reciprocal mechanical cues [83,84]. Thus, in stress relaxing tissues, there is an interplay between strain stiffening through collagen fiber alignment and stress relaxation through collagen fiber reorganization. Studies have shown that through the loss of active cell contraction, collagen tissues relax by about 65%, suggesting that 35% of stress on tissues were relaxed during cell-induced strains [85]. This is corroborated by stress relaxation tests we performed on mouse lung tissue (**Fig. S5**). Thus, in the model, when contractile myofibroblasts die, the stress increase caused by strain-stiffening relaxes by only 65%, in which the remaining 35% of stress is assumed to be relaxed through collagen rearrangement during cell contraction.

**Table 2:** Simulation of ECM behavior.

| Behavior | References |
| --- | --- |
| Baseline stiffness initialized per pixel as a random value between 1 and 3 kPa | Brown, et al., 2013; Liu, et al., 2010 |
| TGFβ1 statically bound to ECM when deposited | Barcellos-Hoff, 1996; Wipff et al., 2007 |
| Collagen content linearly increases ECM stiffness within a given pixel through the formula: $E \text{ (Pa)} = 10 * C_{\text{collagen}} \text{ (mg/mL)}$ | Yang and Kaufman, 2009; Willits and Skornia, 2012 |
| ECM stiffness is increased logarithmically via contraction-mediated strain stiffening through the formula:<br>$E_{\text{traction}} \text{ (kPa)} = 0.5 * \ln(E_{\text{matrix}} \text{ (kPa)})$ | Marinkovic et al., 2012a; Marinkovic et al., 2012b |

#### Collagen agent behavior

While fibroblasts deposit a variety of ECM components, we have chosen to model only type I collagen deposition since it is the most abundant ECM component, it is widely recognized as contributing to ECM stiffening in fibrosis, and it is frequently quantified in fibrosis models to guide model parameterization. A primary role of fibroblasts is to produce ECM, and the rate of collagen deposition changes with the extent of myofibroblast differentiation. Quiescent fibroblasts secrete a small amount of collagen, 15 pg/hr or 1 unit per time step, and proto-myofibroblasts produce collagen at a 5-fold greater rate than quiescent fibroblasts, 75.6 pg/hr or 5 units per time step [86–88]. Collagen does not diffuse in the model but contributes to the stiffness of the matrix in its deposited location. The matrix stiffness increases linearly with collagen concentration, as is described above in the section on ECM mechanical behavior. Collagen crosslinking via lysyl oxidase is excluded from this model due to the lack of available data regarding both the relationship of individual enzyme activity to matrix stiffness as well as how this relationship would change with increasing collagen density and contraction. Collagen turnover is dictated by MMPs and tissue inhibitors of metalloproteinases (TIMPs) [69,89], and an increase in collagen turnover is a characteristic of progressive fibrosis [88,90,91]. While MMPs and TIMPs are not explicitly represented in this model, the influence of these enzymes is simplified into a single parameter that represents daily collagen turnover. At every 24 hour timestep, a random selection of collagen is removed that comprises 7% of the total deposited collagen [92]. The total amount of deposited collagen is tracked over time.

**Table 3:** Collagen agent behavior.

| Behavior | References |
| --- | --- |
| Stiffens matrix proportionally to concentration (See **Fig. S3**) | Marinkovic et al., 2012a; Marinkovic et al., 2012b |
| 7% of randomly selected collagens are degraded daily by a combination of MMP and TIMP activity | Laurent 1987 |
| Relaxes stress in < 30 minutes | Figure S5 |

#### TGFβ1 agent behavior

We focused on the role of TGFβ1 due to its widely recognized role as a profibrotic agent [93–95], it’s unique interactions with the ECM [96], and the fact that it has been shown to be sufficient for activating fibroblasts along the myofibroblast differentiation lineage either alone or in combination with stiffness [97,98]. The model is initialized with a random distribution of TGFβ1 which we assumed to have been released by macrophages in response to tissue damage. TGFβ1 does not diffuse when initially deposited, as it is assumed to be synthesized in the latent form, which binds to the ECM and renders it inactive until it is released by cellular traction forces[99]. Thus, TGFβ1 moves randomly once a fibroblast moves within 10 µm of where it is bound. Additionally, traction forces exerted by contractile myofibroblasts within 50 µm of TGFβ1 will allow TGFβ1 to move randomly, as this is the radius of high strains exerted by contractile myofibroblasts [74]. Non-senescent fibroblasts within 10 µm of bound or unbound TGFβ1 will consume the cytokine. Additionally, proto-myofibroblasts secrete TGFβ1 in an autocrine signaling loop [100]. Specifically, every 7 days 60 pM TGFβ1 per 3,200 cells is released [98], equating to 1 unit of TGFβ1 for every 4 cells in this model.

**Table 4:** TGFβ*1* agent behavior.

| Behavior | References |
| --- | --- |
| Deposited randomly at model initialization | *estimated* |
| Bound to matrix when deposited | Barcellos-Hoff 1996; Wipff et al., 2007 |
| Released from matrix when a cell moves within 10 μm radius or when within 50 μm radius of myofibroblast contraction | Wipff et al., 2007 |

#### Quiescent fibroblast agent behavior

The model is populated with quiescent fibroblasts at initialization that are randomly dispersed throughout the interstitial space. Quiescent fibroblasts migrate within the interstitial space via stiffness-weighted directionality that mimics cellular durotaxis, or directional migration in response to mechanical gradients [101]. Specifically, every 5^th^ time-step, cells evaluate the stiffness of the interstitial tissue regions within 20 µm of the cell and move in the direction of the highest stiffness tissue. Quiescent fibroblasts deposit 15 pg of collagen per hour in the ECM, or 1 unit per time step [102]. When a quiescent fibroblast encounters TGFβ1 bound to the matrix, it both releases that cytokine and consumes it. Quiescent fibroblasts may also consume unbound TGFβ1 that has already been released from the latency complex. For every 1 ng/mL, or one unit, of TGFβ1 a quiescent fibroblast consumes, it replicates once [103]. Once a quiescent fibroblast consumes 5 units of TGFβ1, equivalent to 5 ng/mL, it will differentiate into a proto-myofibroblast [23,104]. Additionally, if a quiescent fibroblast migrates to a region of the ECM with a stiffness greater than 10 kPa it will differentiate into a proto-myofibroblast [23]. Quiescent fibroblasts do not undergo apoptosis.

**Table 5:** Quiescent fibroblast agent behavior.

| Behavior | References |
| --- | --- |
| Migrates in the direction of high stiffness within interstitial space at 10 $\mu\text{m/hr}$ | Lo, et al., 2000; Sunyer, et al., 2020; Espina, et al., 2022 |
| Secretes collagen at 15 pg/hr | Yurovsky, 2002; Islam et al., 2023 |
| Releases TGFβ1 from LAP complex within a radius of 10 $\mu\text{m}$ | Wipff, et al., 2007 |
| Consumes TGFβ1 released from LAP complex within a radius of 10 $\mu\text{m}$ | Shvartsman, et al., 2001; Wipff, et al., 2007 |
| Replicates once after consuming 1 ng/mL of TGFβ1 | Fine and Goldstein, 1987 |
| Differentiates into proto-myofibroblast when exposed to ECM stiffness greater than 10 kPa <u>or</u> 5 ng/mL of TGFβ1 | Hinz, 2006; Hinz, 2010; Klingberg, et al., 2014 |

#### Proto-myofibroblast agent behavior

Proto-myofibroblasts are cells in a transient state of myofibroblast activation. In this model, proto-myofibroblasts arise from quiescent fibroblasts and can either revert to quiescence or die following their lifespan. At the end of the cell’s life span there is one that dies for every 15 proto-myofibroblasts that revert to quiescence [105]. Proto-myofibroblasts synthesize 5 times as much collagen as quiescent fibroblasts, 75.6 pg/hr or 5 units for every time step [86–88], and migrate via stiffness-weighted durotaxis similar to quiescent fibroblasts. Proto-myofibroblasts replicate once upon the differentiation from quiescent fibroblasts. Like quiescent fibroblasts, when a proto-myofibroblast migrates over TGFβ1 that is bound to the matrix, it both releases and consumes the cytokine. Proto-myofibroblasts may also consume unbound TGFβ1. Proto-myofibroblasts must both consume 5 ng/mL of TGFβ1 and co-localize with ECM of greater than 10 kPa stiffness for at least 72 hours to differentiate into contractile myofibroblasts [78,106,107].

**Table 6:** Proto-myofibroblast agent behavior.

| Behavior | References |
| --- | --- |
| Migrates in the direction of high stiffness within interstitial space at 10 $\mu\text{m/hr}$ | Lo et al., 2000; Trichet et al., 2012; Shi et al., 2015 |
| Secretes collagen at 75.6 pg/hr (5x quiescent fibroblasts) | Burgess et al., 2005; Jones et al., 2010; Hansen et al., 2016 |
| Replicates once upon transition from quiescent to proto-myofibroblast | Grotendorst et al., 2004; Wynn and Ramalingam, 2012 |
| Releases TGFβ1 from LAP complex within a radius of 10 $\mu\text{m}$ | Wipff et al., 2007 |
| Consumes TGFβ1 released from LAP complex within a radius of 10 $\mu\text{m}$ | Shvartsman et al., 2001 |
| Differentiates into myofibroblast when exposed to 2 ng/mL of TGFβ1 <u>and</u> ECM stiffness greater than 10 kPa for 72 hours | Hinz et al., 2001; Marinkovic et al., 2012; Shi et al., 2013 |
| Reverts to quiescence or undergoes apoptosis following its lifetime of 72 hours at a ratio of 15:1 | Kisseleva et al., 2012 |

#### Contractile myofibroblast behavior

Contractile myofibroblasts represent fully activated myofibroblasts with αSMA^+^ stress fibers and exert high levels of contractile forces on the matrix. Contractile myofibroblasts arise only from proto-myofibroblasts after an extended period of pro-fibrotic cues, as described in the previous section. Contractile myofibroblasts do not migrate, but exhibit high levels of traction on their local ECM [106]. This traction-mediated ECM strain is assumed to be uniform on all interstitial tissue within a radius of 100 µm [74,106,108]. This cellular traction induces strain stiffening in the ECM in a manner dependent on the stiffness of the tissue at each pixel as discussed in the section on ECM mechanics above. Contractile myofibroblasts may still uptake TGFβ1. After a lifespan of 10 days, contractile myofibroblasts die through apoptosis [109,110]. Contractile myofibroblasts do not revert to quiescence or transient proto-myofibroblasts.

**Table 7:** Contractile myofibroblast agent behavior.

| Behavior | References |
| --- | --- |
| Contracts ECM within a radius of 100 $\mu\text{m}$ (See **Fig. S4**) | Hinz et al., 2001; Han et al., 2018; Pakshir et al. 2019 |
| Remains stationary | Thampatty and Wang, 2006; Younesi et al., 2021 |
| Undergoes apoptosis after 10 days | Desmouliere et al., 1995; Darby, et al., 1990 |

### Sensitivity Analysis

Model parameters were evaluated for their influence on collagen deposition and tissue stiffness model outputs. Each parameter was varied by an increase of 10% and 20%, as well as a decrease of 10% and 20%. Each parameter set was simulated in 5 replicates. Collagen deposition and tissue stiffness were independently plotted against the input parameter and a two-tailed correlation analysis was performed with 𝛼 = 0.05 as the significance cutoff. Model sensitivity is shown in **Fig. S6**.

### Model Validation

The young mouse and aged mouse ABM outputs were validated against experimental data gathered from both our own mouse studies as well as dozens of published studies. Data from published studies was extracted from graphs using WebPlotDigitizer (automeris.io) and mean values ± SD were plotted against model outputs mean values ± SD. All values were converted to reflect values per lung (all lobes) for a male mouse. Model outputs were extrapolated assuming uniformity across the lung, and assuming a lung volume of 0.075 cm^3^.

#### Collagen Validation

Collagen secretion values were taken from studies that used the bleomycin mouse model of lung fibrosis and assessed collagen using the hydroxyproline assay. Hydroxyproline was assumed to make up 13% of collagen proteins on average, and values were converted to micrograms (µg) per lung.

#### Tissue Mechanics Validation

Reported and experimentally derived mechanical data were used to validate model outputs. Literature values used in the validation were limited to atomic force microscopy (AFM) data of lung tissue from either the bleomycin mouse model or IPF patients. These values were compared to the overall stiffness over time of the simulation area. Additionally, stiffness maps were exported at the end of the simulation and compared with nanoindentation data generated by our mouse studies, described above. Raw simulation data were scaled to more closely resemble the length scale of the experimental data. To do this, five 10 µm patches across and five 10 µm patches down were averaged into a single point to simulate the 50 µm indentation probe. This scaled mechanical map was then used for subsequent comparisons.

Simulated and experimentally-derived mechanical maps were quantitatively compared to assess variability and the potential influence on pulmonary function. The potential influence on lung function was assessed by measuring the stiffness burden, defined as the percent of the measured area that was above 20 kPa. These values were each compared using an unpaired t-test.

#### TGFβ1 Validation

TGFβ1 accumulation data was derived from papers that used ELISA kits to evaluate total TGFβ1 concentration from their lung tissue homogenate. TGFβ1 measurements taken on bronchoalveolar lavage (BAL) fluid was discarded as this is not the tissue compartment that we are modeling and concentrations are difficult to normalize between studies. In the model, TGFβ1 was categorized into active and inactive depending on whether it was bound to the matrix or not. All TGFβ1 that the model is initialized with is assumed to be in its inactive state and bound to the matrix.

### Aged Mouse model adaptation

Following the validation of the young mouse model, parameters known to change with age were altered for subsequent analysis. A summary of parameter adaptations made to fit the aged mouse mode as listed in **Table 8**. Additionally, the model was initialized using histology images gathered from the lungs of mice aged to 21 months.

**Table 8:**
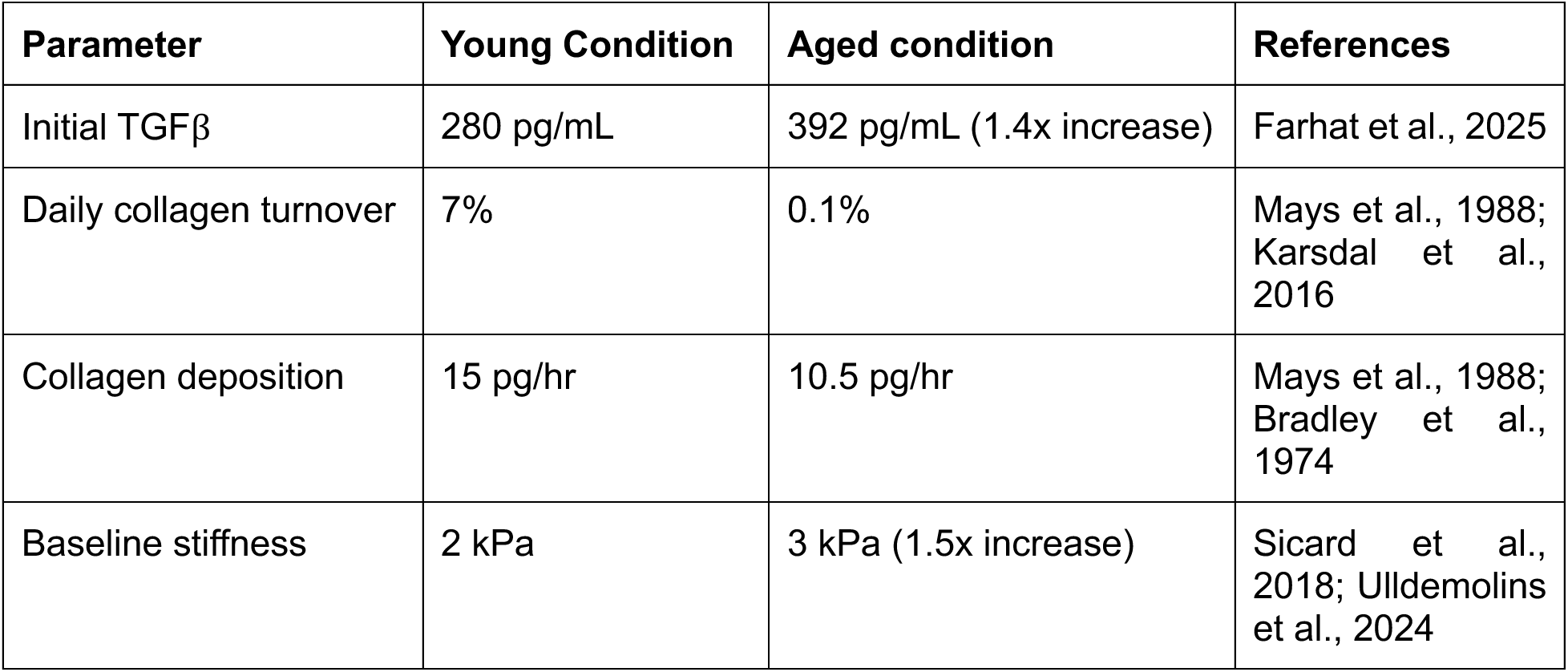
Age-dependent model parameters.

### Senescent cell model

#### Senescent cell behavior

At the initialization of the model, 3 to 15 senescent fibroblasts are randomly distributed within the interstitial space of the simulation area, along with quiescent fibroblasts and TGF𝛽1 cytokines. Senescent fibroblasts migrate at 4 µm/hr in a random manner [111], and secrete low levels of collagen at a rate of 5 pg/hr [112,113]. Further, senescent cells exert low levels of contractile forces on the ECM (**Fig. S4**) [114]. As is characteristic of senescent cells, they do not undergo replication [115]. Non-senescent fibroblasts may become senescent through direct contact with senescent cells, assumed to be mediated by the cytoplasmic transfer of reactive oxygen species [116], although additional studies demonstrated that NOTCH-mediated cell-cell contact may also transmit senescence [117,118]. Finally, contractile myofibroblasts may become senescent at the end of their lifespan as a means of apoptosis evasion [25]. A summary of senescent cell behaviors is provided in **Table 9**.

#### Static senescent cell simulations

To investigate the influence of senescent cell behavior in a controlled manner, we held the number of senescent fibroblasts constant in each simulation. Thus, we initialized 3 to 15 senescent fibroblasts at the start of the simulation and removed all other sources of senescent fibroblasts in the model. Specifically, we removed juxtacrine-mediated secondary senescence and the differentiation of contractile myofibroblasts into senescence fibroblasts. Additionally, we varied the lifespan of these senescent fibroblasts, and removed the immune-mediated clearance of senescent cells. The amount of TGFβ1 secreted by senescent fibroblasts, their lifespan, and the number of senescent fibroblasts at the start of the simulation were varied in a full factorial. The identity and range of these parameters are outline below in **Table 10**, as well as the baseline levels derived from literature. Model simulations were run to 14 days with 5 replicate model runs per parameter set.

**Table 9:**
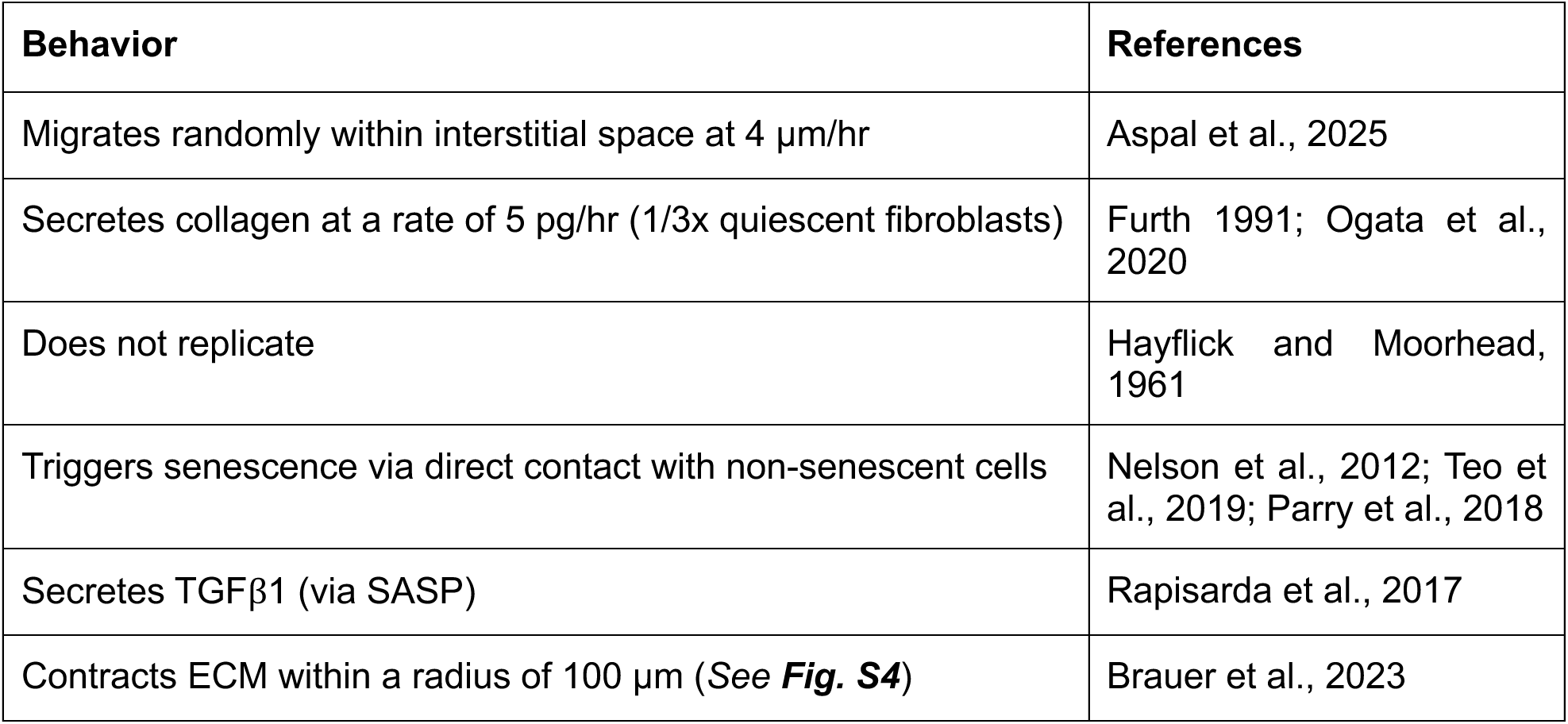
Senescent fibroblast agent behavior.

| Behavior | References |
| --- | --- |
| Migrates randomly within interstitial space at 4 $\mu\text{m/hr}$ | Aspal et al., 2025 |
| Secretes collagen at a rate of 5 pg/hr (1/3x quiescent fibroblasts) | Furth 1991; Ogata et al., 2020 |
| Does not replicate | Hayflick and Moorhead, 1961 |
| Triggers senescence via direct contact with non-senescent cells | Nelson et al., 2012; Teo et al., 2019; Parry et al., 2018 |
| Secretes TGFβ1 (via SASP) | Rapisarda et al., 2017 |
| Contracts ECM within a radius of 100 $\mu\text{m}$ (See **Fig. S4**) | Brauer et al., 2023 |

**Table 10:** Static senescence simulation parameter ranges.

| Parameter | Range | Baseline level | References |
| --- | --- | --- | --- |
| TGFβ1 secretion (TGFb-amount) | 0, 10, 20, 40 ng/hr | 20 ng/hr | Basisty et al., 2020; Rapisarda et al., 2017 |
| Senescent cell lifespan (senescent-lifespan) | 5 days, 10 days, 14 days | 6 days | *estimated* |
| Number of senescent cells present at model initialization (sen-fibroblast-count) | 3 cells, 6 cells, 9 cells, 12 cells, 15 cells | 3 cells | Anerillas et al., 2023 |

#### Dynamic senescent cell simulations

To investigate the parameters defining interactions between senescent fibroblasts and other cell types for their relative contribution to fibrosis, parameters were estimated and varied across the ranges outlined in **Table 11**. Specifically, the time required for senescence to transmit via cell-cell contact (ROS-transfer-time) and the percentage of contractile myofibroblasts that become senescent at the end of their lifespan (p-senescence) were each varied in full factorial while holding all other parameters static, including the elimination of immune-mediated senescent cell clearance. Additionally, the two parameters that dictate immune cell clearance were varied across their range while holding the remaining parameters at their estimated baseline levels. These parameters include the percentage of senescent cells cleared every 24 hours (p-clearance), and the lag in immune cell entry before daily clearance begins (immune-cell-entrance). Model simulations were run to 14 days with 5 replicate model runs per parameter set.

**Table 11:** Senescent cell dynamics simulated parameter ranges.

| Parameter | Range | Baseline level | References |
| --- | --- | --- | --- |
| Time required for juxtacrine-mediated secondary senescence (ROS-transfer-time) | 30 min, 60 min, 90 min, 2 hr, 2.5 hr | 1 hour | Nelson et al., 2012 |
| Probability of contractile myofibroblasts that become senescent at the end of their lifespan (p-senescence) | 10%, 20%, 30%, 40%, 50% | 20% | Jun and Lau, 2010 |
| Probability that senescent cells are eliminated by daily immune infiltration (p-clearance) | 0, 10% 25%, 75%, 100% | 50% | Chandrasegaran et al., 2025 |
| Time delay of immune cell infiltration (immune-cell-entrance) | 0, 3, 5, 7, 9 days | 3 days | *estimated* |

## Results

### Young mouse model outputs align with experimental trends and values

We utilized the large number of studies leveraging the bleomycin model of lung fibrosis in young mice to validate emergent behavior within our model. Fibroblast collagen secretion in the model was compared against reported hydroxyproline data, and both values were converted to reflect the excess amount of collagen per lung, compared to the non-fibrotic control. While there is some discrepancy in the range of collagen deposition between published studies, the model output aligns well with both the general trend of increasing collagen over time, as well as the values reported from multiple independent studies [58,119–125] (**Fig. 2a**). Tissue stiffness generated from the model was compared to indentation measurements of both human IPF patient and fibrotic mouse lung tissue, from five independent studies [71,126–129] as well as our own. Simulated tissue stiffness aligns well with literature results, where day 14 and 21 stiffness values are more in line with mouse lung stiffnesses than human values (**Fig. 2b**). Bound and active TGFβ1 amounts from the model were compared to reported values of active TGFβ1 from tissue digestion measured via ELISA. Model TGFβ1 concentration follows the same trend as literature [130,131], albeit at lower concentrations (**Fig. 2c**). This is likely because the only ongoing source of TGFβ1 secretion in the model is autocrine signaling of fibroblasts, whereas in biological systems there are many more cell types that contribute to available TGFβ1 [132]. Together, these data show this model to be an accurate, but conservative, estimate of the progression of fibrosis in the lung.

**Figure 2:**
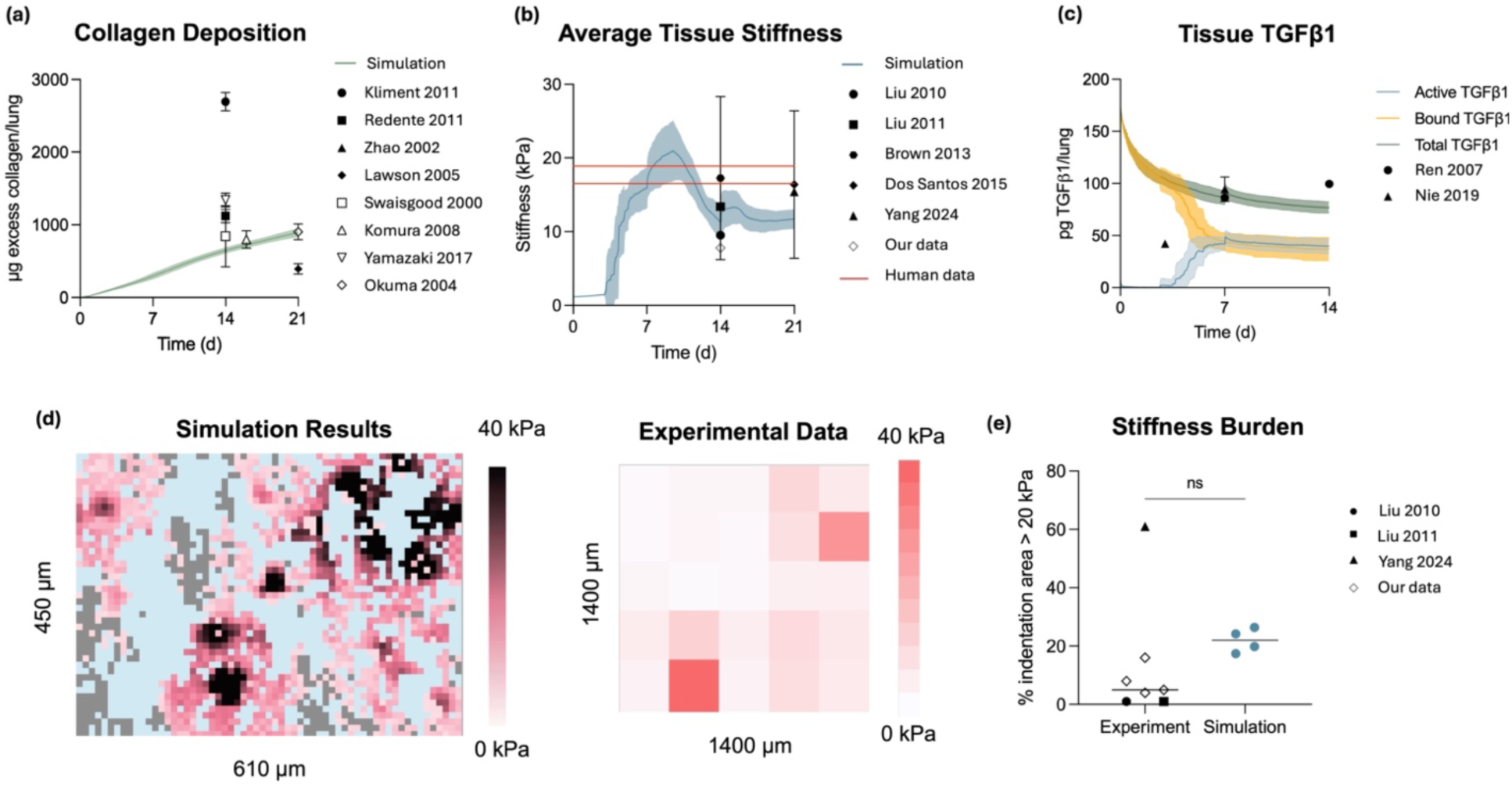
Young mouse model validation. In (a) – (c) all simulated values are plotted as lines with shading corresponding to SD. These simulations correspond to one input histology image. Experimentally-derived values are plotted with points +/- SD. **(a)** Simulated collagen deposition amounts fall within the range of experimentally-derived values. **(b)** Simulated tissue stiffness values fall within the range of experimentally-derived measurements of mouse lung stiffness following bleomycin exposure. Human data derived from Booth et al., 2012 and Hilster et al., 2020. **(c)** Simulated active TGFβ1 amounts (unbound from latency complex) follow the same trend as experimentally-derived values. **(d)** Representative images of simulated and experimentally-derived mechanical maps at day 14 post-bleomycin installation. **(e)** Quantification of the stiffness burden from experimentally-derived or simulated mechanical maps. Each point represents one mouse. For each mouse, 5 replicate simulations were run per histology image with 2 unique histology images. These 10 simulations were averaged. Groups compared via unpaired t-test with Welch’s correction.

Further, we compared mechanical maps generated by nanoindentation of bleomycin-exposed mouse lungs to simulated mechanical maps generated by the model. Representative examples of simulated and experimentally derived mechanical maps are shown in **Fig. 2d**. From these mechanical maps we quantified the stiffness burden, defined as the percentage of the indentation area measuring > 20 kPa, to gain a general sense of the influence of scar tissue buildup on lung function. Despite the simulated mechanical maps encompassing a significantly smaller area than our experimentally-derived maps and a considerably larger area than literature-derived AFM maps, no significant difference in mechanical burden was observed between the experimental and simulated mouse lungs (**Fig. 2e**).

### Aged mouse model captures high stiffness burden in fibrotic lung

Following validation of the young mouse model, we then adjusted input parameters to mimic fibroblast behaviors that change with age (**Table 8**). These variations include changing lung architecture, captured via the input of aged control mouse lung histology, an increase in the TGFβ1 present at the start of the simulation [133], a decrease in collagen turnover [134,135] and deposition [134,136], and an increase in starting ECM stiffness [137,138]. This aged mouse model was then validated against available literature data and experimentally-derived information regarding aged mouse fibrotic lungs. Simulated collagen deposition, again, followed the same increasing trend as literature values [58,139–141] and fell within the range of values reported for each timepoint (**Fig. 3a**). Additionally, simulated mouse lung stiffness aligned with two unique animal models of lung fibrosis in aged mice, as well as with the stiffness of human IPF lungs (**Fig. 3b**). There was no literature data available for tissue-derived TGFβ1 in aged mouse lung, although studies report a 2- to 20-fold increase in TGFβ1 mRNA expression in aged mice compared to young mice [133,139]. Our simulations show an order of magnitude increase in TGFβ1 expression (**Fig. 3c**), which falls in line with the range of reported increase.

**Figure 3:**
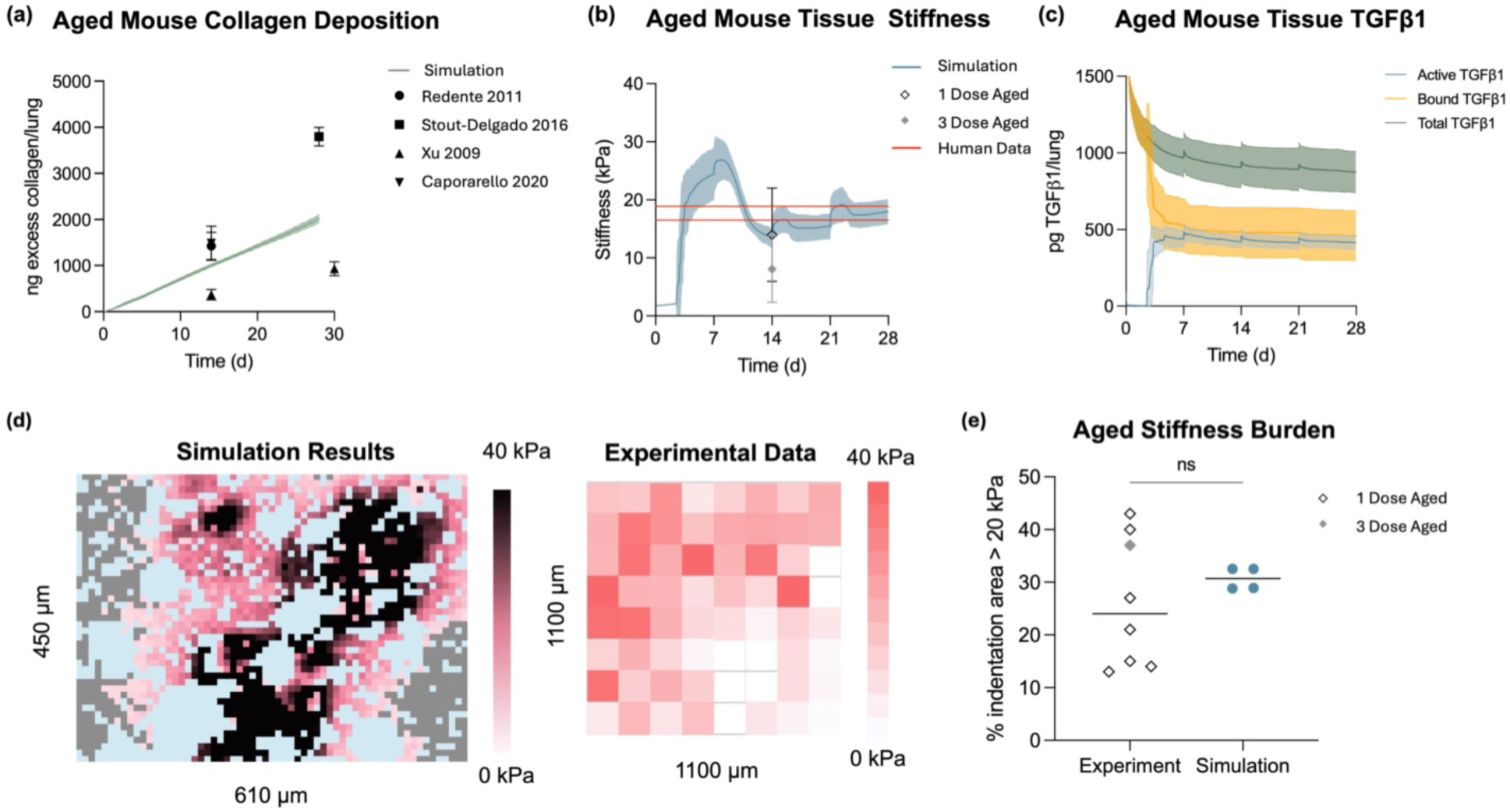
Aged mouse model validation. In (a) and (b) simulated values are plotted as lines with shading corresponding to SD. These simulations correspond to one input histology image. Experimentally-derived values are plotted with a point +/- SD. **(a)** Simulated collagen deposition falls in the range of experimentally-derived values from aged mice. **(b)** Simulated tissue stiffness falls within the range of stiffness values determined from two animal models, as well as human data. Human data derived from Booth et al., 2012 and Hilster et al., 2020. **(c)** Aged mouse TGFβ1 amounts follow the same trend as those of young mice. **(d)** Representative images of simulated and experimentally-derived mechanical maps at day 14 post-bleomycin installation. Experimental data is derived from three-dose aged mouse lung. **(e)** Quantification of the stiffness burden from experimentally-derived or simulated mechanical maps. Each point represents one mouse. For each mouse, 5 replicate simulations were run per histology image with 2 unique histology images. These 10 simulations were averaged. Groups compared via unpaired t-test with Welch’s correction.

Changes in the stiffness burden of aged mouse lung was also assessed by comparing mechanical maps generated through nanoindentation of bleomycin-dosed aged mice. Representative mechanical maps are shown in **Fig. 3d**, and the % area of these maps greater than 20 kPa is quantified in **Fig. 3e**. Both one-dose and three-dose aged mice show remarkable alignment with the simulated stiffness burden from the aged mouse model, demonstrating the robustness of this simulation.

### An increase in static senescent cell burden contributes significantly to fibrosis in the lung

Following validation of the aged mouse model, we then layered on senescent cell behaviors in stages to investigate their relative influence (**Fig. 4**). First, we sought to examine how the presence of senescent cells in isolation contributes to fibrosis progression. To do this, we held the number of senescent cells in the simulation constant by removing interactions between agents that change the number of senescent cells present, including secondary senescence, apoptosis evasion, and immune cell clearance. Then we varied the parameters that control the senescent cell burden, the amount of TGFβ1 secreted by senescent cells, and the persistence of senescent cells (**Table 10**).

**Figure 4:**
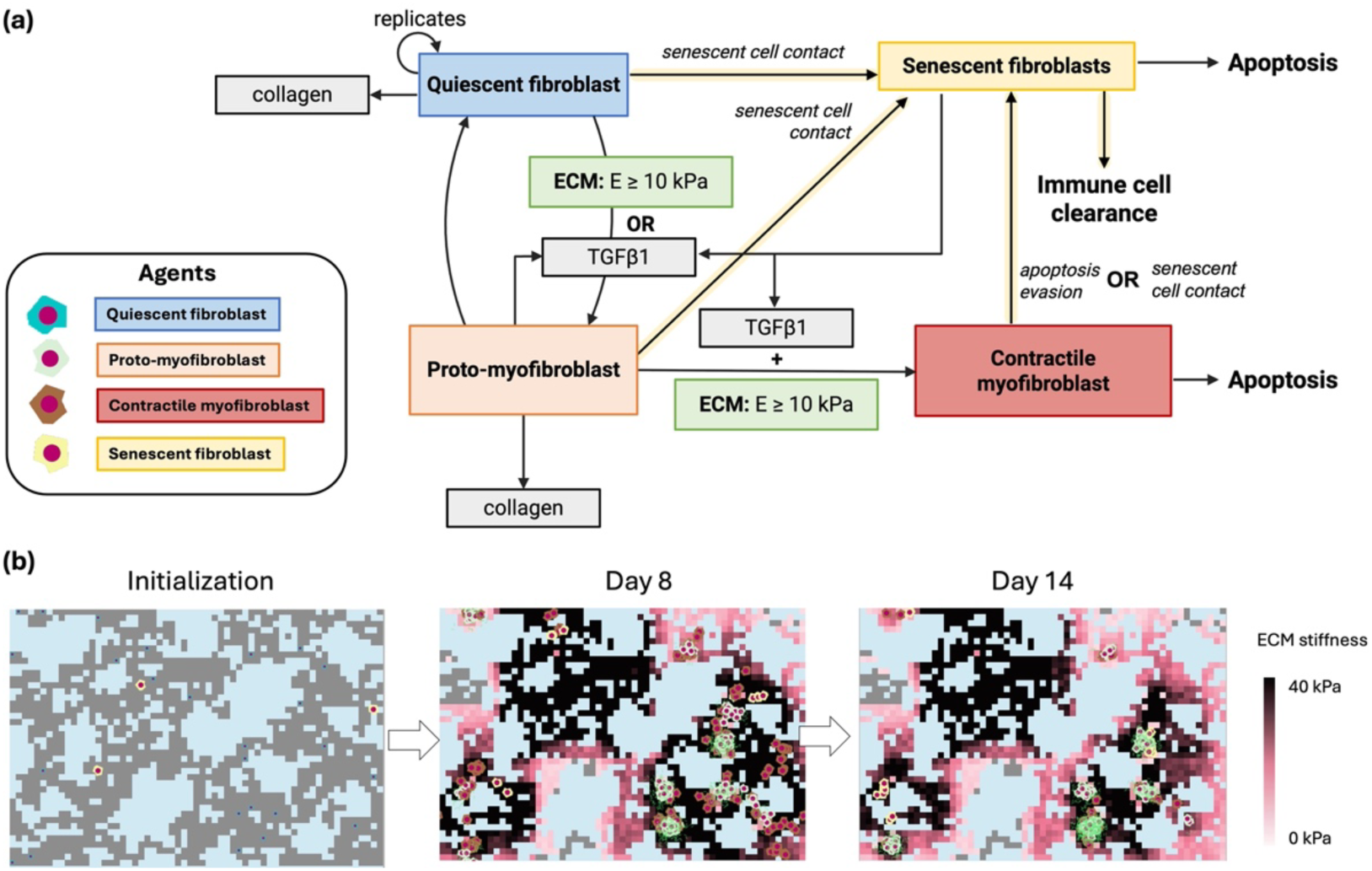
Senescent fibroblast behavior was incorporated into the aged mouse model. **(a)** Senescent fibroblast agents were incorporated into existing pathways in the model as depicted in the schematic. Specifically, senescent fibroblasts secrete TGFβ1 as a part of their secretome, trigger secondary senescence in any nearby cell type through direct contact, and are removed by assumed immune cell intervention. Additionally, contractile myofibroblasts may evade apoptosis through becoming senescent and senescent cells themselves eventually apoptosis. Lines highlighted in yellow denote interactions that are removed during static simulations as they govern the total number of senescent cells. **(b)** Example images represent how the simulation may progress at baseline conditions described in **Tables 10 and 11**.

We first performed a sensitivity analysis on these three parameters to assess their contribution to simulated tissue stiffness and collagen accumulation. There was not a significant correlation between collagen accumulation and the total number or lifespan of senescent cells (p = 0.973 and p = 0.941 respectively, **Fig. 5a**). Tissue stiffness at day 14 showed some level of sensitivity to all three parameters, though it was influenced much less by the persistence of senescent cells than the total number of senescent cells or the TGFβ1 secretion rate of these cells. To assess the relative contributions of the three variables across the full parameter space, we applied multivariate linear regression with either tissue stiffness or collagen accumulation as the dependent variable (**Fig. S7**). We found that these parameters did not exhibit a linear relationship with collagen accumulation, again indicating these parameters have minimal influence on collagen deposition. In contrast, tissue stiffness showed a high degree of linearity in the multivariate regression analysis against these parameters, with the number of senescent cells showing the largest standardized coefficient among the predictors. Further, we assessed the relative influence of the total number of senescent cells and their TGFβ1 secretion rate in the case of senescent cell persistence through the duration of the simulation (14 days). Although both parameters demonstrated a strong influence on simulated tissue stiffness, the influence of the number of senescent cells, again, was more significant than the amount of TGFβ1 that these cells secrete (**Fig. 5b**). As may be expected, these parameters are closely related and we found that the greater the number of senescent cells present, the more influential their TGFb1 secretion rate becomes (**Fig. S8**). Taken together, these data point to the accumulation of senescent cells as a primary contributor toward fibrosis progression in this model.

**Figure 5:**
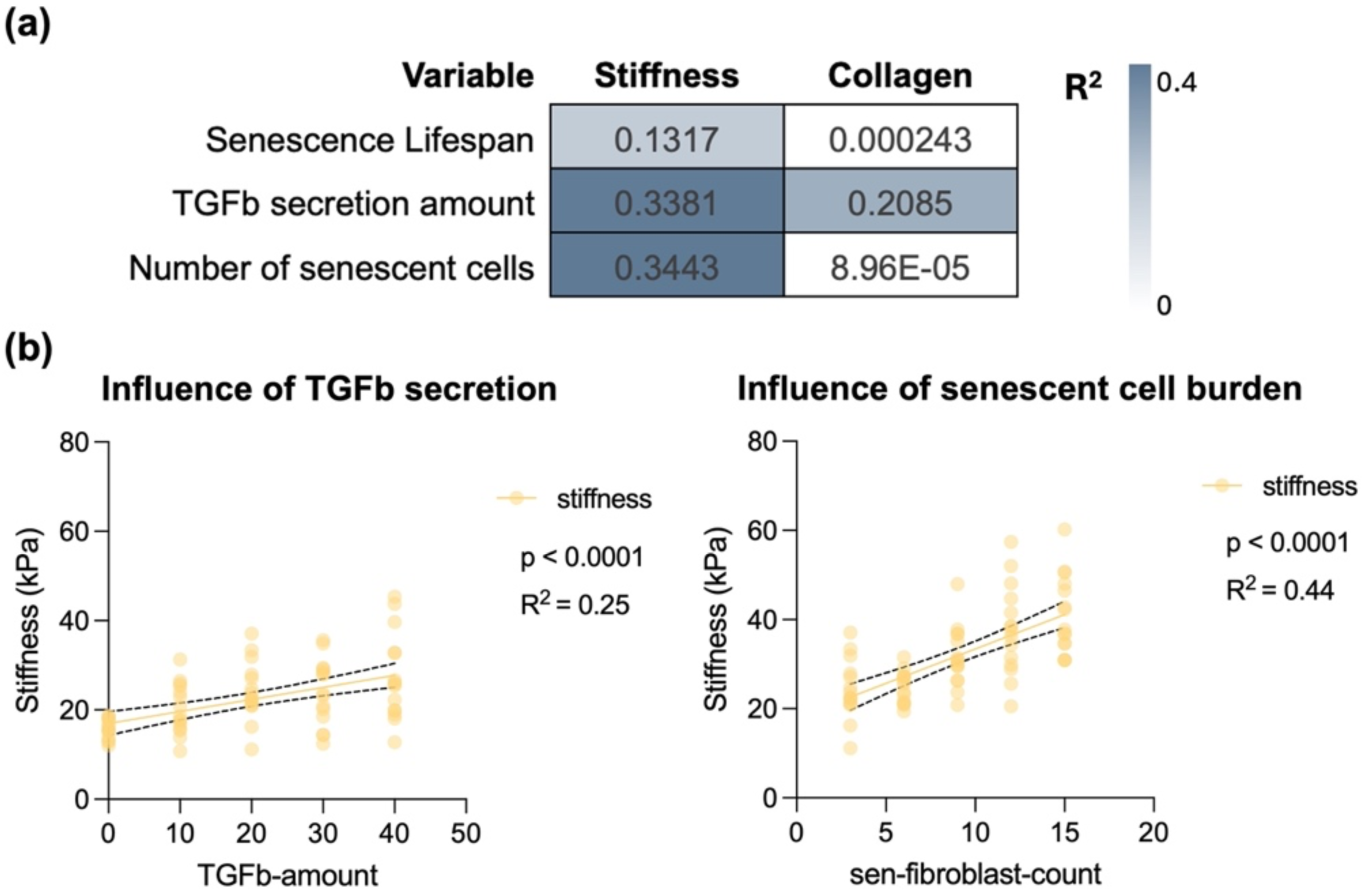
Static senescent cell burden and TGFβ1 secretion rate strongly influence tissue stiffness. **(a)** Sensitivity analysis was conducted by varying each of the three parameters while holding the others at baseline. Correlation analysis was performed and R^2^ values are plotted in the heatmap. Boxes shown in white did not have a significant correlation. **(b)** Senescent cell lifespan was extended to 14 days, and the correlation between either senescent cell number or TGFβ1 secretion was assessed. Each point represents one simulation run across the range of parameters evaluated.

### Senescent cell burden is influenced more by juxtacrine-mediated bystander senescence than apoptosis resistance

We next sought to investigate which components of cell-cell dynamics represented in this model made the strongest contribution to overall senescent cell burden. Specifically, we evaluated the relative influence of juxtacrine-mediated bystander senescence and contractile myofibroblast senescence in the absence of immune cell clearance. We evaluated the dynamics between myofibroblast subtypes over time across the entirety of the parameter space (**Fig. 6a**). We see from the relative consistency in the pattern of these time-sweep graphs across increasing values in the probability of apoptosis-mediated contractile myofibroblast senescence (horizontally across graphs), indicating this parameter has a relatively small effect on fibroblast subtype. On the contrary, we see drastic differences in the pattern of these graphs vertically across changing values in the time required for juxtacrine-mediated secondary senescence, indicating this variable plays a much more prominent role on fibroblast subtype. Looking specifically at the senescent cell burden in these simulations (orange line, yellow shading), we see again that the time required for juxtacrine-mediated secondary senescence plays a dominant role in the number of senescent cells throughout the simulation. For instance, the senescent cell burden on average does not reach over 100 cells per simulation space in simulations where the time required for bystander senescence is at least 2 hours, regardless of contractile myofibroblast senescence probability. Taken together, we can conclude that juxtacrine-mediated secondary senescence plays a much more prominent role in the overall senescent cell burden in these simulations than contractile myofibroblast senescence.

**Figure 6:**
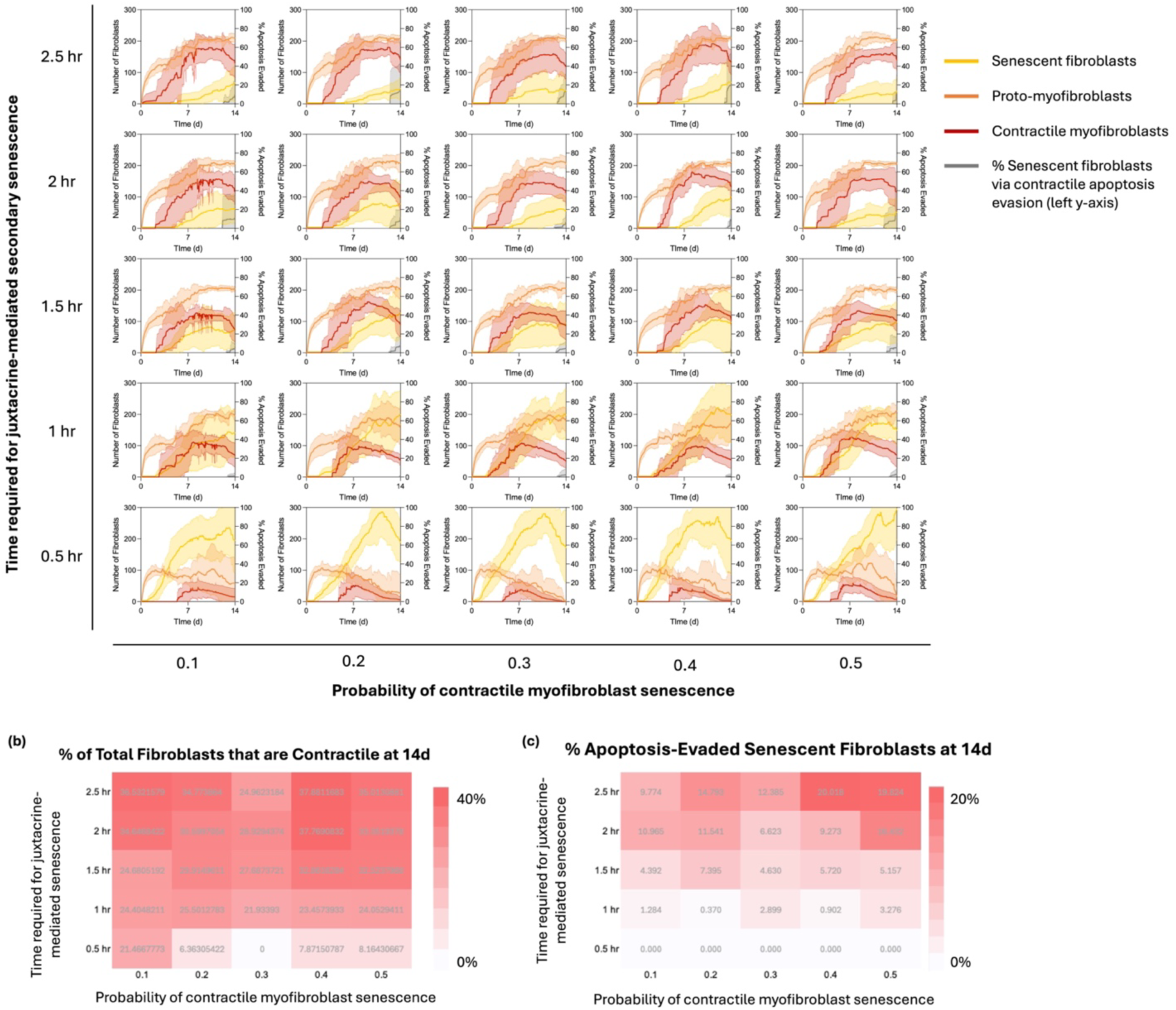
Senescent cell burden is influenced more by juxtacrine-mediated bystander senescence than apoptosis resistance. **(a)** A two-way parameter sweep was performed to investigate the relative contribution of the time-constraint that mediated juxtacrine-mediated senescence and the probability of myofibroblast senescence. The number of fibroblast subtypes as well as the percent of senescent fibroblasts that underwent apoptosis evasion were plotted as a function of time across the simulation. The lines represent an average value across 9 simulations, 3 technical replicates per unique histology image. **(b)** The percent of senescent fibroblasts that arose from contractile myofibroblast evasion of apoptosis at the endpoint of the simulation (14 days) was plotted as a heat map to compare the relative contribution of these two variables. **(c)** The percent of non-senescent fibroblasts (including quiescent fibroblasts) that are contractile at the endpoint of the simulation (14 days) was plotted as a heat map to compare the relative contribution of these two variables.

The strong effect of bystander senescence may be expected in this model, as the overall pool of cells able to undergo senescence is greater than contractile fibroblasts alone. To investigate the relative contribution of this disparity, we considered the number of contractile myofibroblasts relative to the other types of fibroblasts that can become senescent at the endpoint of the simulation (**Fig. 6b**). We saw from this analysis that there is not a strong influence of either of these two parameters on the percentage of contractile fibroblasts. Additionally, we looked at the relative contribution of contractile myofibroblasts that evaded apoptosis to become senescent in the overall pool of senescent fibroblasts (**Fig. 6c**). Here, however, we do see influence of these parameters on the portion of senescent cells that come from apoptosis evasion. The values for the percentage of senescent fibroblasts that have evaded apoptosis are plotted at each time point on the right y-axis of each sub-graph in **Fig. 6a** and the values at 14 days simulation are plotted in the heatmap in **Fig. 6c**. In both cases, it seems there is a combined influence of both parameters. To evaluate this analytically, we again performed multivariate linear regression analysis to assess the relative contribution of these two parameters on the percentage of senescent cells that have evaded apoptosis (**Fig. S9**). We see a high degree of linearity in this analysis, indicating an excellent model fit. Additionally, we see almost identical contributions of these two parameters on the percent of senescent fibroblasts that have evaded apoptosis. Taken together, these results show us that while juxtacrine-mediated secondary senescence has a greater influence on the overall number of senescent cells, these two parameters have equal weight in their relative contribution to the senescent cell pool.

### Timing of immune cell clearance of senescent fibroblasts mediates fibrotic outcome

Finally, we investigated the influence of immune cell dynamics on senescent cell accumulation. The efficiency of immune cell-mediated senescent cell clearance (p-clearance), as well as the timing of immune cell entry, were each varied across their parameter space and assessed according to changes in fibroblast subtypes (**Fig. 7a**). These simulations revealed interesting emergent behavior. As would be expected, for lower levels of senescent cell clearance efficiency, there was an overall greater number of senescent cells over the course of the simulations. Highly efficient clearance of senescent cells from the onset of the simulation led to unchecked myofibroblast accumulation, especially in the case of proto-myofibroblasts (**Fig. 7a**, top left graph). However, for a delay in immune cell entrance of 3 days or more, there is clearance of senescent cells as well as myofibroblast populations by 14 days. This recession of both myofibroblasts and senescent cells to low levels (below ∼ 50 cells) by 14 days was only seen for scenarios of delayed immune cells entrance of 3 days or more and of senescent cell clearance efficiency of 50% or greater (depicted in the green box). Further investigation of these parameters in isolation revealed that the number of senescent cells at day 14 was strongly negatively regulated by the percentage of senescent cells cleared each day, but not influenced by the timing of immune cell entrance (**Fig. 7b**). However, the timing of immune cell delay did have a strong influence on the overall fibrotic burden, as seen by the negative correlation with both simulated collagen accumulation and tissue stiffness (**Fig. 7c**). These correlations were not seen in the percentage of senescent cells cleared (**Fig. 7d**). Thus, these data predict a nuanced role of senescent cell clearance in lung fibrosis progression.

**Figure 7:**
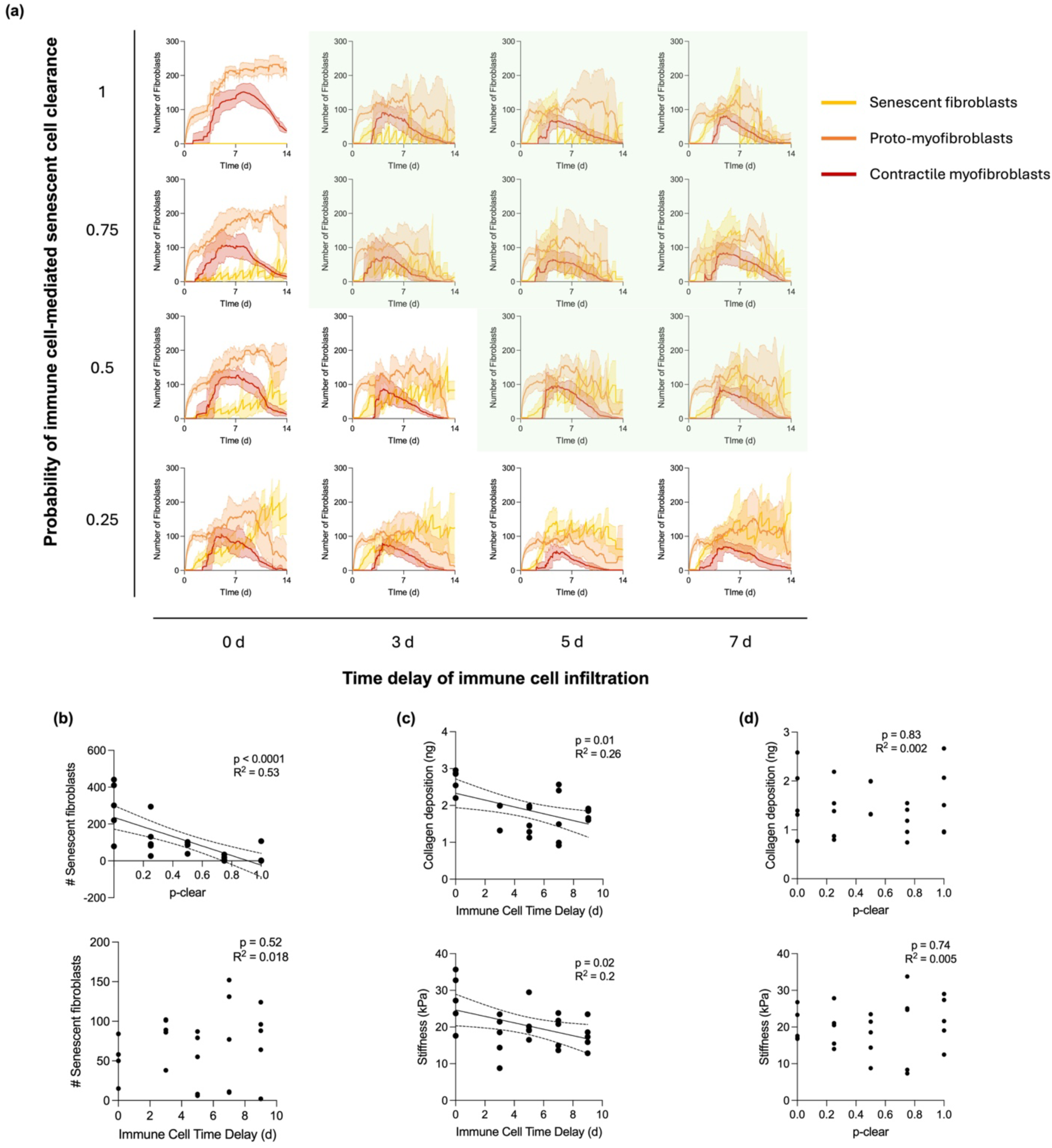
Timing of immune cell clearance of senescent cells mediates fibrotic outcome. **(a)** A two-way parameter sweep was performed to investigate the dynamics of immune-mediated senescent cell clearance. Graphs in the green box indicate parameter combinations that led to < 100 senescent cells and myofibroblasts at 14 days. **(b)** Additionally, correlation analysis was performed for each of these parameters in isolation on the accumulation of senescent cells, **(c)-(d)** as well as on tissue stiffness and collagen accumulation at day 14.

## Discussion

We developed a new ABM that simulates fibrosis in the lung of young and aged mice to investigate how senescent cell dynamics contribute to fibrosis progression. To our knowledge, this is the first computational model to integrate the distinct functional roles of fibroblast subtypes in myofibroblast activation alongside dynamic changes in ECM mechanics during fibrosis progression. Simulated changes in tissue stiffness and collagen accumulation aligned well with experimental data for both young and aged mouse models. Thus, our aged mouse ABM provided a controlled and informative test bed for investigating the influence of senescent fibroblast behavior in fibrosis progression.

Through our simulations, we found that in the absence of immune cell regulation, the overall number of senescent cells has the largest impact on fibrosis progression, and this senescent cell burden is strongly regulated by bystander senescence induction, or senescence that is triggered by nearby senescent cells. Furthermore, the model’s emergent behavior demonstrated that the timing and efficiency of immune-mediated senescent cell clearance are co-dependent factors required for proper wound healing. These insights provide grounds for future experimental investigation and warrant consideration into the timing of senolytics treatment for IPF.

This model enabled unique predictions regarding the changes in lung mechanics with fibroblast differentiation. As is commonly reported, we observed the reproducible emergence of fibroblast-rich foci, accompanied by high levels of ECM stiffness and collagen deposition. Additionally, we noted that sudden increases in tissue stiffness during simulations coincide regularly with activation of contractile myofibroblasts. This suggests that strain stiffening contributes a much greater portion to tissue stiffness than collagen deposition. We verified this hypothesis by evaluating changes in tissue stiffness in the absence of strain-stiffening behavior, and observed a drastic reduction in tissue stiffness (**Fig. S10**). The co-occurrence of contractile myofibroblast differentiation and an increase in contraction-mediated tissue stiffness explains the high levels of tissue stiffness that we see around 7-10 days, that then recedes by day 14 as these contractile myofibroblasts die (**Fig. 2b, Fig. 3b)**. While others have postulated this dominant role of strain stiffening in IPF [68], the disregard of collagen crosslinking in this model may be undermining the true contribution of collagen in tissue stiffness. Thus, additional experimental data investigating the relationship between enzyme-mediated collagen crosslinking and contracted ECM on lung tissue stiffness would offer important clarity into the driving force behind this pathological stiffening.

Additionally, recent evidence has shown that fibroblast durotaxis may be a driver of fibrosis progression in *in vivo* models of lung fibrosis [142]. In this model, durotaxis was simulated using stiffness-weighted directional migration at every n^th^ time step, and the frequency of this durotactic migration was calibrated as a part of model validation. This calibration revealed novel insights regarding the relationship between stiffness-mediated migration and fibrotic architecture. Namely, the greater the influence of durotactic migration (and subsequent decrease in random migration) the more tightly compact the high-stiffness foci became (**Fig. S11**). With less frequent durotactic migration the regions of high stiffness were more diffuse, leading to a general increase in tissue stiffness. These results corroborate previous findings that *in vivo* fibrotic lung tissues exhibit steep stiffness gradients, and further implicate durotactic behavior in establishing these gradients.

Regarding senescent cell behavior, this model demonstrated bystander senescence as a primary contributor to overall senescent cell burden. These results may be exacerbated by the exclusion of paracrine-mediated senescence, although there is a lack of experimental consensus as to whether secondary senescence is transmitted via juxtacrine signaling, paracrine signaling, or both. Given the lack of experimental data describing secondary senescence, we chose to simulate only juxtacrine-mediated secondary senescence, as this seemed to be the most conservative approach with the fewest required parameter estimations. Further, a recent model of senescence in wound healing also demonstrated the dominant role of juxtacrine-mediated senescence, even in the presence of paracrine-mediated secondary senescence [57]. Taken together, these data emphasize the need to further investigate the mechanisms that regulate secondary senescence, given its prominent contribution to the senescent cell population.

Additionally, the model’s emergent behavior regarding the influence of immune clearance of senescent cells offers useful insights into their role in tissue remodeling. A previous model of senescent cell behavior in dermal wound healing revealed the presence of pre-existing senescent cells in a wound led to persistent inflammation, whereas the absence of pre-existing senescent cells in a wound led to transient inflammation [57]. We observed similar results in our lung-specific model. Namely, the immediate clearance of senescent cells removed any negative regulation on the myofibroblast population, leading to a greater degree of tissue fibrosis. Conversely, the inefficient clearance of the senescent cell population led to an over-expansion of senescent cells in the simulation, which may cause persistent inflammation (not simulated). This highlights the importance of coordinated senescent cell appearance and clearance during wound healing, as has been discussed elsewhere [143–145]. These data may offer useful insights into study design and treatment approaches for ongoing clinical trials investigating senolytics for IPF treatment.

While this model shows remarkable agreement with many of the experimental validation metrics, there are many key characteristics of lung fibrosis that are not represented in our model, providing avenues for future investigation. Specifically, repeated injury to the lung epithelium is considered the initiating event in pulmonary fibrosis, and epithelial cell dysregulation is known to contribute significantly to fibrosis progression [146–148]. Thus, incorporating epithelial cell behavior into this model may yield interesting new insights.

Additionally, although simplified immune regulation is represented in this model, the distinct cell types and varied phenotypes of immune cells are largely overlooked. Given the importance of immune regulation on fibrosis progression [149–151], the implication of these complexities in the immune compartment are vital to understand. Further, these cell types are each regulated by a host of cytokines beyond the pro-fibrotic influence of TGFβ1, and interactions between these cytokines are important to model to more fully capture the complexities of the *in vivo* microenvironment. While this model operates effectively with relatively few rules and agents, the introduction of these relevant interactions may enable investigation of more complex and nuanced biological questions.

Additionally, this model assumes a simplified and monotonic senescent cell phenotype, opening the opportunity for deeper investigation into the multifaceted influence of senescent cells in lung fibrosis. First, studies have shown that senescent cell phenotype varies significantly with both the cell type that is induced to senescence [152–155] and the means by which it is induced [156,157]. Additionally, epithelial cell senescence has been shown to be an important contributor to the overall senescent burden in IPF [9]. Thus, the incorporation of distinct senescent cell phenotypes would improve model accuracy. Further, the SASP is a characteristic component of senescent cell phenotype that was highly simplified in this model. Future model iterations may improve upon this representation with a more thorough representation of SASP components including its direct link to immune cell infiltration. Overall, our model provides useful insights into the general influence of senescent cells on fibrosis in the lung, and future work may expand upon this to further investigate other known regulators of the fibrotic milieu.

## Conclusions

This work describes the development of a novel ABM for the investigation of senescent cell behavior in pulmonary fibrosis progression. By leveraging experimental data from both young and aged mouse lungs, we were able to construct a model of fibrosis progression with outputs that align well with reported values. Through the incorporation of senescent cell behaviors into the aged mouse model, we were able to investigate unique characteristics of senescent cell interactions in a layered fashion to assess their role in fibrosis progression. These simulations revealed the importance of juxtacrine-mediated secondary senescence in its contribution to the senescent cell population, as well as the complex interplay between senescent cell accumulation and immune-clearance in mediating tissue outcomes. Overall, this model provides useful insights into the dynamics of senescent fibroblasts in pulmonary fibrosis that may be further evaluated experimentally.

## Acknowledgments

We would like to thank Dr. Jenna Sumey for her assistance with young and aged mouse lung mechanical characterization, as well as Sheri VanHoose from UVA’s Research Histology Core for all her assistance in processing, sectioning and staining of mouse lung tissue. This work was supported by the NIH (R35GM138187, R35GM162024, R01HL179312, F32HL170760 to R.T.H) and NSF (CAREER DMR/BMAT 2046592, GRFP to M.L.S.). The content is solely the responsibility of the authors and does not necessarily represent the official views of the National Institutes of Health.

