## Supporting Information for "An agent-based model suggests how senescent cell behavior and matrix mechanics drive pulmonary fibrosis in aged mice"

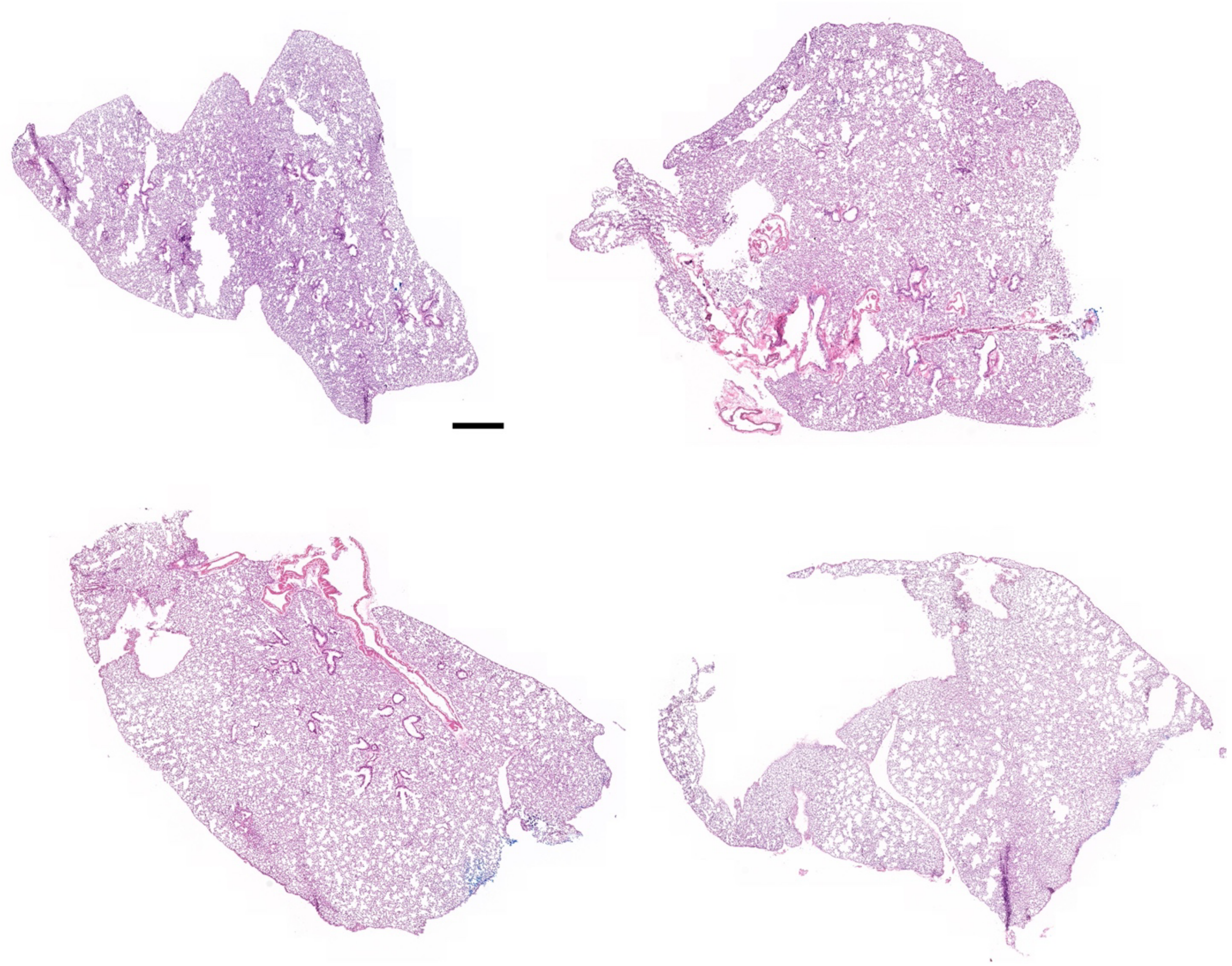

**Figure S1:** Whole-lung histology images from 14-week-old control mice. Scale bar: 500 µm.

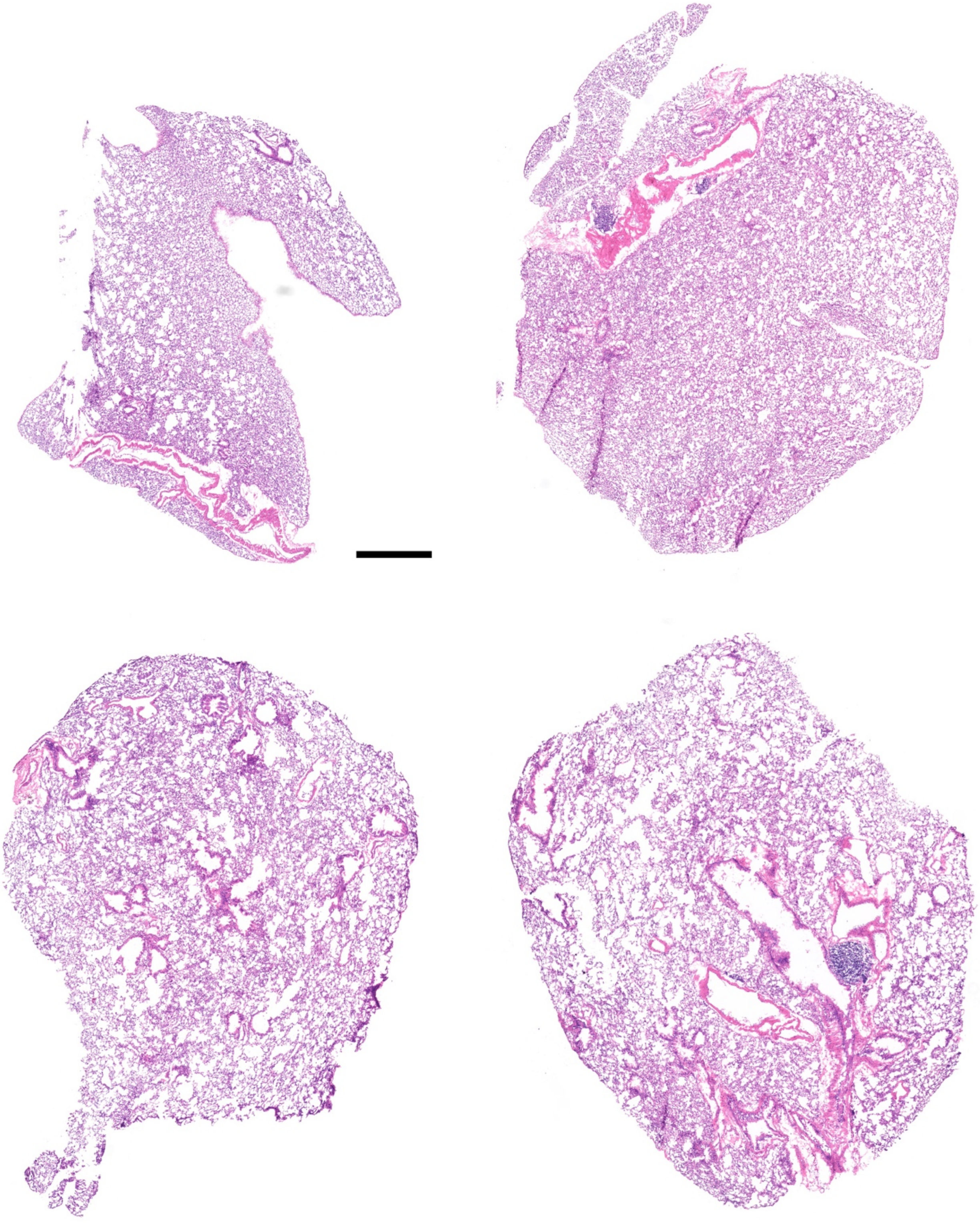

**Figure S2:** Whole-lung histology images from 16-month-old control mice. Scale bar: 500 µm.

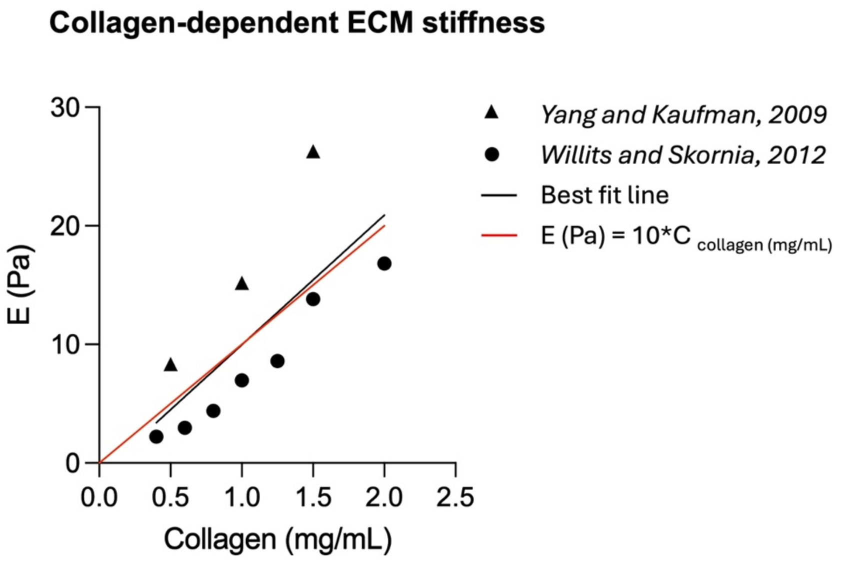


**Figure S3:** Collagen gel stiffness was plotted against the concentration of collagen used to determine the relationship between collagen content and ECM stiffness. These data were derived from two independent sources and fit to a linear model. The line of best fit is shown in black. The linear model was simplified to the line shown in red and implemented in the ABM.

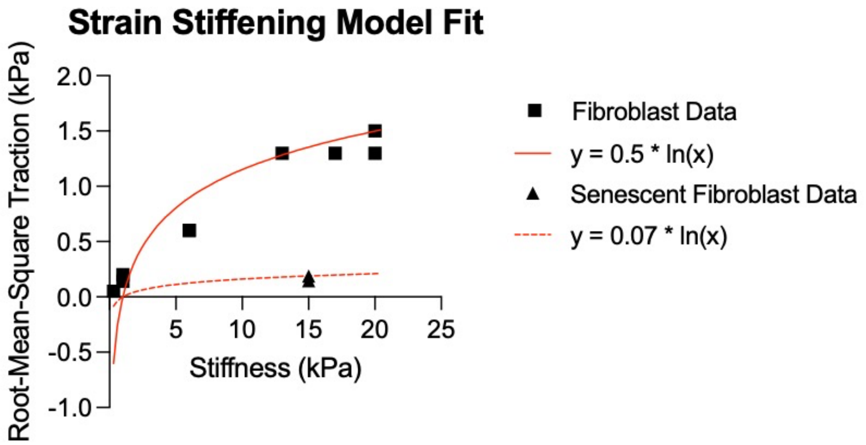


**Figure S4:** Traction force microscopy data from two independent sources (Marinkovic et al., 2012a and Marinkovic et al., 2012b). Fibroblast traction force data were plotted against the stiffness of the hydrogel cell culture substrate. These data were fit to a logarithmic model as shown by the solid red line. Additionally, senescent cell traction force data were derived from Brauer et al., 2023 and the logarithmic model was scaled to fit the data, as shown by the dashed red line.

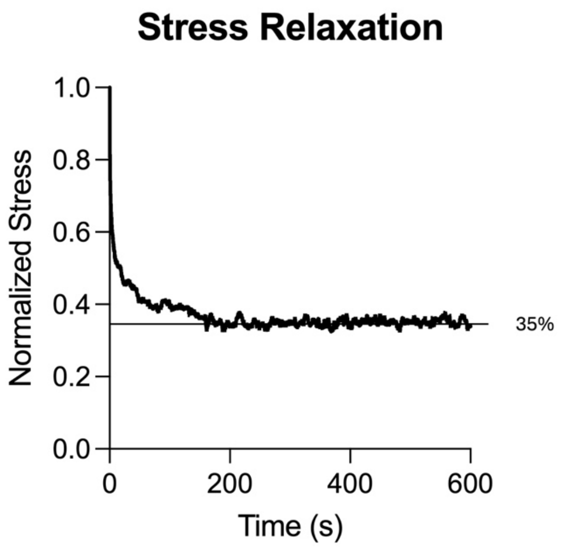


**Figure S5:** Young mouse lung indentation was used to assess stress relaxation over an extended time. All stress values were normalized to the maximum stress and plotted over time. Normalized stress levels out to about 35% of the maximum stress.

**(a) Correlation Significance Summary**

| Variable | Stiffness | Collagen |
|---|---|---|
| Fibroblast Lifespan | | |
| Contraction Radius | + , ** | |
| Fibroblast Count | | + , * |
| Mechanical Activation | | - , * |
| Myofibroblast Lifespan | | |
| Collagen Deposition | | + , * |
| Collagen Turnover | | - , ** |
| Contractile Lifespan | + , * | |
| Cytokine Count | | + , ** |
| n-step | +, ** | |
| Reversion Rate | | |

**(b) Correlation Strength Summary**

| Variable | Stiffness | Collagen |
|---|---|---|
| Fibroblast Lifespan | 0.3066 | 0.1076 |
| Contraction Radius | 0.9235 | 0.07992 |
| Fibroblast Count | 0.1522 | 0.8654 |
| Mechanical Activation | 0.7482 | 0.9098 |
| Myofibroblast Lifespan | 0.00002975 | 0.5621 |
| Collagen Deposition | 0.6783 | 0.8642 |
| Collagen Turnover | 0.6119 | 0.9557 |
| Contractile Lifespan | 0.8879 | 0.07584 |
| Cytokine Count | 0.3203 | 0.9262 |
| n-step | 0.9223 | 0.4103 |
| Reversion Rate | 0.5565 | 0.009367 |

**Figure S6:** Sensitivity analysis was performed for all relevant parameters. Parameters were varied up 20% in 10% increments (+20%, +10%) and down 20% in 10% increments (-20%, -10%). Tissue stiffness and collagen accumulation were plotted against the input variable and two-tailed correlation analysis was performed. **(a)** Parameters that the model are not significantly correlated with are denoted with grey boxes. Parameters that have a significant effect on model outcomes are denoted with either + or - to indicate whether they are positively or negatively correlated, and an asterisk to denote the level of significance. *p < 0.0332, **p < 0.0021. **(b)** $R^2$ values are plotted in the heatmap, in which the darker blue indicates a higher $R^2$. Values are listed in the boxes in grey.

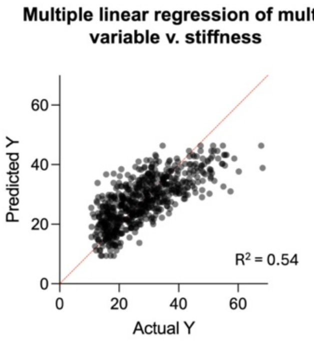


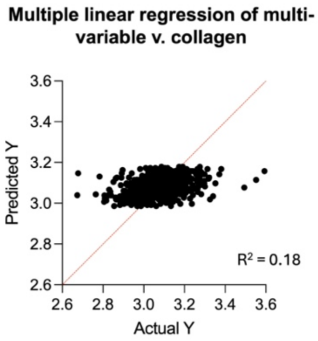


**Stiffness Multiple Linear Regression Model Summary**

| Parameter estimates | Variable | Estimate | Standard error | 95% CI (asymptotic) | \|t\| | P value | P value summary |
|---|---|---|---|---|---|---|---|
| β0 | Intercept | 4.528 | 1.021 | 2.522 to 6.534 | 4.434 | | |
| β1 | TGFb-amount | 0.4325 | 0.02085 | 0.3916 to 0.4734 | 20.75 | <0.0001 | **** |
| β2 | sen-fibroblast-count | 1.044 | 0.06949 | 0.9071 to 1.180 | 15.02 | <0.0001 | **** |
| β3 | senescent-lifespan | 0.007401 | 0.0008686 | 0.005695 to 0.009107 | 8.520 | <0.0001 | **** |

**Collagen Multiple Linear Regression Model Summary**

| Parameter estimates | Variable | Estimate | Standard error | 95% CI (asymptotic) | \|t\| | P value | P value summary |
|---|---|---|---|---|---|---|---|
| β0 | Intercept | 3.002 | 0.01442 | 2.974 to 3.030 | 208.3 | | |
| β1 | TGFb-amount | 0.003234 | 0.0002942 | 0.002656 to 0.003812 | 10.99 | <0.0001 | **** |
| β2 | sen-fibroblast-count | 0.003494 | 0.0009808 | 0.001568 to 0.005420 | 3.563 | 0.0004 | *** |
| β3 | senescent-lifespan | -2.169e-005 | 1.226e-005 | -4.577e-005 to 2.386e-006 | 1.769 | 0.0774 | ns |

**Figure S7:** A multivariate regression model was fit to the static senescence model output data to assess which parameter related to senescent cell behavior had the greatest influence on fibrotic phenotype. The input variables dictating the amount of TGFβ1 senescent cells secrete (TGFb-amount), the total number of senescent cells present (sen-fibroblast-count), and the senescent cell lifespan (senescent-lifespan) were evaluated against two output variables – tissue stiffness and collagen deposition. The graph on the top left refers to the model in which tissue stiffness is the response variable, and this model seems to reflect a high degree of linearity, meaning tissue stiffness is responsive to changes in these input variables. The first table describes the parameters of the model fit for the stiffness model, and comparison of the β parameter estimates shows the number of senescent fibroblasts to have the greatest influence on tissue stiffness. Additionally, the graph on the top right shows the model in which collagen deposition is the response variable, and this model does not show a high degree of linearity, meaning this linear regression analysis is not an appropriate method for distinguishing trends in this dataset. The summary of parameters of the collagen model is shown in the bottom table.

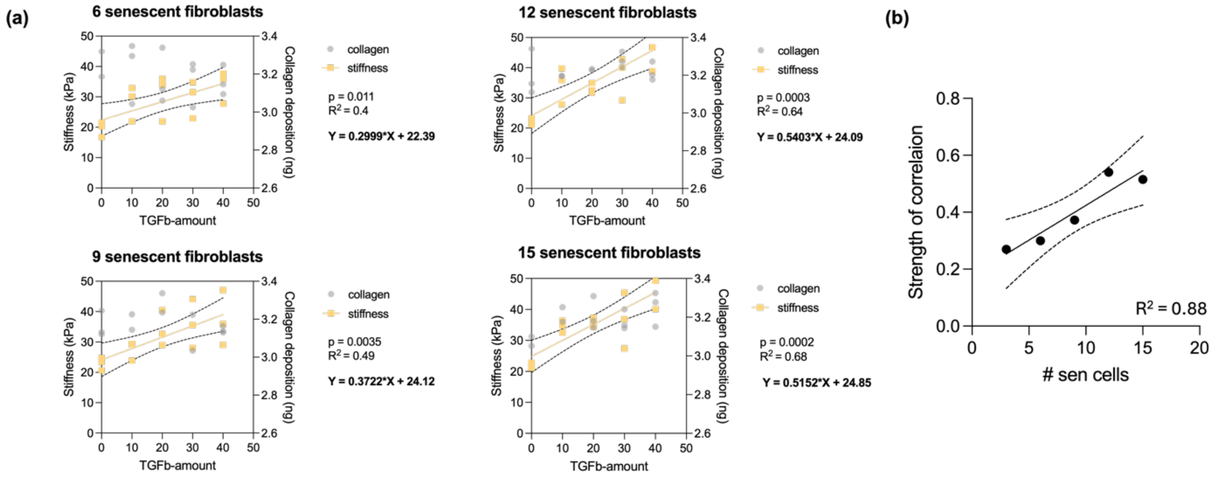


**Figure S8:** In the static senescent cell simulation, the amount of TGFβ1 secreted is correlated with increased tissue stiffness, and this correlation is sensitive to the number of senescent cells present. **(a)** The tissue stiffness and collagen accumulation for three replicates of each parameter set are shown. Each graph represents a different number of senescent cells present, and the x-axis denotes the amount of TGFβ1 secreted per cell in ng/hour. Each of these graphs shows a linear correlation with tissue stiffness, but not with collagen deposition. The p-value, $R^2$ value, and equation for each linear fit is shown to the right of each graph. **(b)** To assess the relationship between the number of senescent cells present and the influence of TGFβ1 secretion on final tissue stiffness, we plotted the slope of each correlation as shown in the graphs in (a) as a function of the number of senescent cells simulated. This graph also follows a linear trend, with an $R^2$ value of 0.88, indicating that the influence of the amount of TGFβ1 secreted by each senescent cell on the final tissue stiffness is sensitive to the number of senescent cells present.

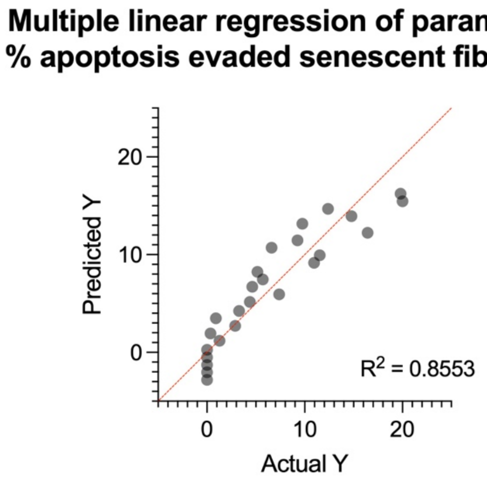


**Model Summary**

| Parameter estimates | Variable | Estimate | 95% CI (profile likelihood) | \|t\| | P value | P value summary |
|---|---|---|---|---|---|---|
| β0 | Intercept | -7.577 | -10.88 to -4.269 | 4.75 | <0.0001 | **** |
| β1 | p-senescence | 7.988 | 6.508 to 9.467 | 11.2 | <0.0001 | **** |
| β2 | ROS-transfer-time | 7.672 | 0.2753 to 15.07 | 2.151 | 0.0427 | * |

**Figure S9:** To assess the relative influence of the two parameters explored in **Fig. 6** on senescent cell burden, we again performed multiple linear regression analysis. The input parameters included the probability of contractile myofibroblasts becoming senescent at the end of their lifetime (p-senescence) and the time required for juxtacrine-mediated secondary senescence of any non-senescent fibroblast (ROS-transfer time). The output variable of this model was the percentage of total senescent fibroblasts that became senescent through contractile myofibroblast apoptosis evasion (as dictated by the first parameter) at the endpoint of the simulation (14 days). This model exhibits a high degree of linearity with an $R^2$ value of 0.855. Comparing the influence of each of these input parameters on the output variable, β values are almost identical, indicating that these two parameters have similar influence on the percentage of senescent cells that underwent apoptosis evasion.

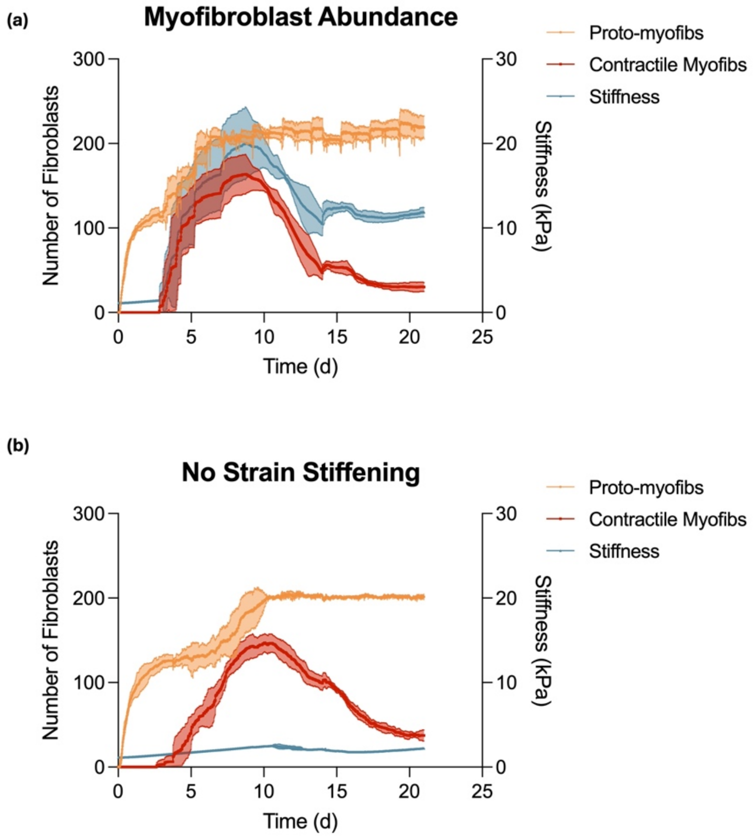


**Figure S10:** To assess the relative influence of contractile myofibroblast-mediated strain stiffening on tissue stiffness, we plotted myofibroblast abundance (left y-axis) and tissue stiffness (right y-axis) as a function of time. Each line represents the mean of 9 simulations and the shaded regions around the lines correspond to the standard deviation. **(a)** In the normal fibrotic simulation, we observe a rapid spike in the abundance of contractile myofibroblasts (shown in red) that coincides with a rapid spike in tissue stiffness (shown in blue). This suggests these model components may be correlated. **(b)** We removed the strain stiffening behavior from contractile myofibroblasts and again ran the simulation. We observe the same pattern of myofibroblast abundance, but there is no spike in tissue stiffness in these simulations. The final tissue stiffness in these simulations remains similar to the starting value, whereas in the presence of stain stiffening (a) tissue stiffness increases significantly.

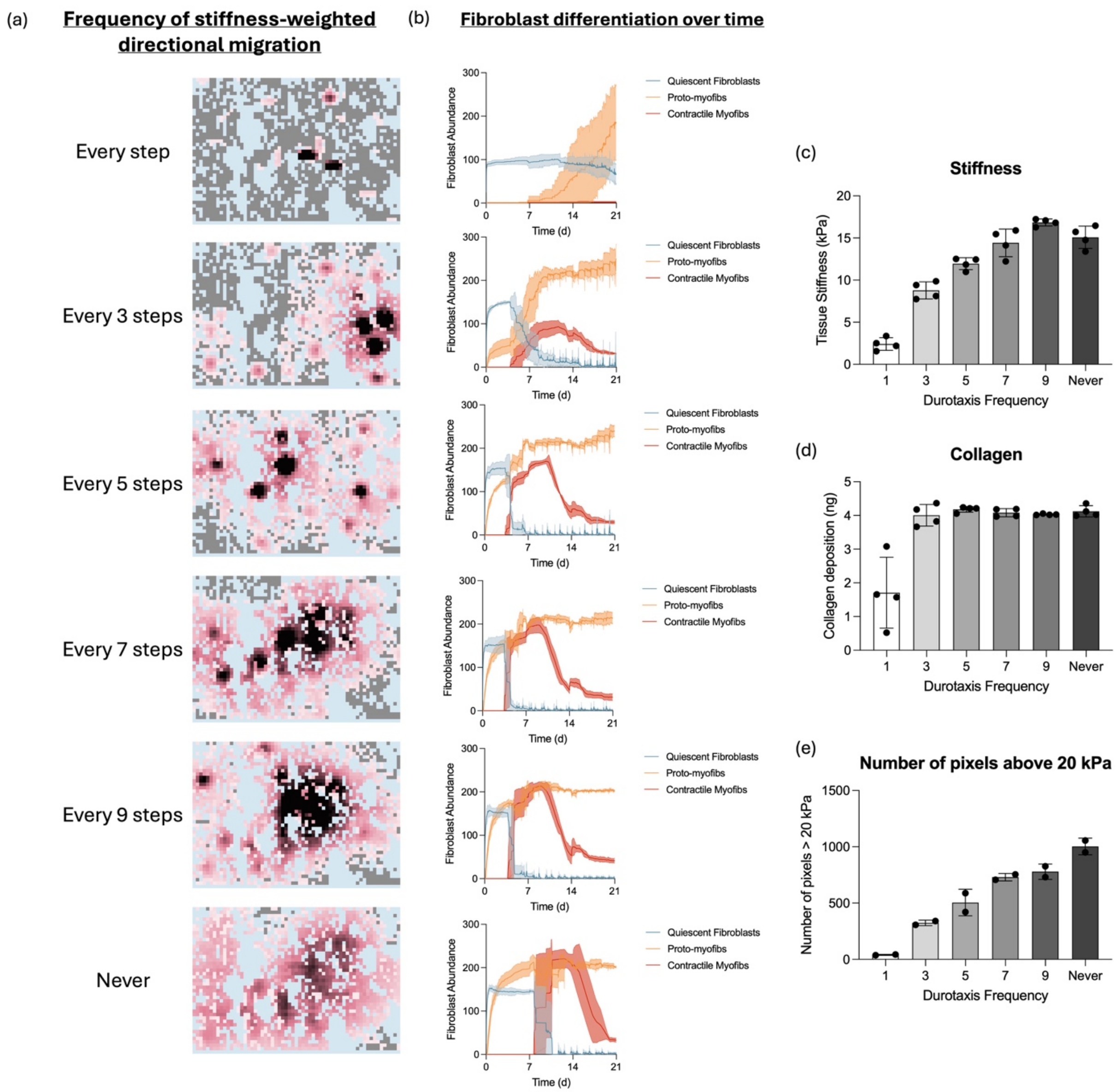


**Figure S11: (a)** Representative images are shown from the endpoint of simulations (21 days) in which the frequency of stiffness-weighted directional migration (durotactic behavior) was varied. Blue pixels represent airspace, grey pixels represent tissue, and pink pixels represent increasing stiffness with darkening hues. **(b)** The abundance of fibroblast subtypes over time was plotted in which each graph again corresponds to a different frequency of durotactic behavior. Durotaxis at every step leads to no contractile myofibroblast differentiation and spots of concentrated tissue stiffness. Increasing the randomness of fibroblast migration enables the full differentiation of contractile myofibroblasts and leads to more diffuse regions of high tissue stiffness. For these simulations, we plotted the final tissue stiffness **(c)** and collagen accumulation **(d)** as a function of durotaxis frequency. Both model outputs increase with greater randomness up to a threshold. **(e)** Additionally, to evaluate the total stiffness burden, we plotted the number of pixels above 20 kPa and saw that this also increases as a function of increasing randomness.
